# Structural Insights into Biased Signaling at Chemokine Receptor CCR7

**DOI:** 10.1101/2025.11.24.690334

**Authors:** Kotaro Tanaka, Kouki Nishikawa, Yuki Shiimura, Yoshinori Fujiyoshi, Naotaka Tsutsumi

**Affiliations:** Cellular and Structural Physiology Laboratory, Institute of Integrated Research, Institute of Science Tokyo, Bunkyo, Tokyo 113-8510, Japan; Joint Research Course for Advanced Biomolecular Characterization, Faculty of Agriculture, Tokyo University of Agriculture and Technology, Fuchu, Tokyo 183-8509, Japan; Division of Molecular Genetics, Institute of Life Science, Kurume University, Fukuoka 830-0011, Japan; Department of Cell Biology, Graduate School of Medicine, Kyoto University, Kyoto 606-8501, Japan

**Author notes:** **Corresponding author:** Naotaka Tsutsumi, **Email:**. **Author contributions:** KT: Data Curation (equal), Formal Analysis (lead), Investigation (supporting), Methodology (equal), Software (lead), Validation (equal), Visualization (equal), Writing – Original Draft (supporting), Writing – Review & Editing (equal). KN: Data Curation (supporting), Investigation (equal), Writing – Review & Editing (supporting). YS: Data Curation (supporting), Formal Analysis (supporting), Investigation (equal), Methodology (equal), Visualization (supporting), Writing – Review & Editing (equal). YF: Funding acquisition (lead), Methodology (equal), Resource (lead), Supervision (equal), Writing – Review & Editing (supporting). NT: Conceptualization (lead), Data Curation (equal), Formal Analysis (supporting), Funding Acquisition (supporting), Investigation (lead), Methodology (equal), Project Administration (lead), Resources (supporting), Software (supporting), Supervision (equal), Validation (equal), Visualization (equal), Writing – Original Draft (lead), Writing – Review & Editing (lead). **Competing interests:** The authors declare no competing interests.

**Keywords:** CCR7, Chemokine, Cryo-EM, MD Simulations, Signaling Bias

## Abstract

CC chemokine receptor 7 (CCR7), which orchestrates adaptive immunity, exhibits a phenomenon known as biased agonism. CCL19 induces robust G protein signaling and β-arrestin recruitment, leading to transient signaling. In contrast, CCL21 preferentially activates G protein pathways with minimal arrestin engagement, resulting in sustained signaling and differential functional outcomes. Here, we present the cryo-EM structures of the human CCR7-G_i_ complex with either CCL19 or CCL21. The structures reveal that while both engage a conserved orthosteric pocket, they adopt markedly distinct binding poses. Notably, the compact 30s loop of CCL21 inserts deeply into the receptor’s extracellular vestibule, whereas the corresponding loop of CCL19 rests atop extracellular loop 2. Molecular dynamics simulations further reveal that these distinct binding modes induce differential intracellular dynamics, linked to the rotameric state of Y83 at the intracellular end of transmembrane helix 1. We demonstrate that CCL19 stabilizes a flexible conformational ensemble with a highly dynamic helix 8, creating a lateral opening favorable for GPCR kinase engagement. Conversely, CCL21 restricts this flexibility, locking the receptor in a state that precludes kinase interaction while maintaining G protein coupling. Corroborated by functional data, these findings provide key insights into the structural basis of biased agonism at CCR7 and establish a foundation for rational design of pathway-selective immunomodulators.

**Significance statement:** Chemokine receptor CCR7 directs immune cell migration by activating distinct signaling pathways in response to CCL19 and CCL21, a phenomenon known as biased agonism. The structural basis for this divergence has remained unclear. By combining cryo-electron microscopy with molecular dynamics simulations, we reveal that although both ligands stabilize similar G protein-bound conformations, they drive distinct receptor dynamics. We demonstrate that CCL19 induces a flexible intracellular conformation necessary for recruiting regulatory kinases, whereas CCL21 locks the receptor in a state precluding this interaction. This “dynamic selection” model aligns with functional data and rationalizes a fundamental paradox in chemokine biology, highlighting the critical role of protein dynamics in signal transduction and providing a blueprint for designing precision immunotherapeutics.

## Introduction

Adaptive immunity relies on the precise spatiotemporal orchestration of immune cell positioning and activation to mount effective responses against pathogens and malignant cells while maintaining self-tolerance (1, 2). Central to this process is CC chemokine receptor 7 (CCR7) (SI Appendix, Fig. S1A), a G protein-coupled receptor (GPCR) that governs the trafficking of various T cells, mature dendritic cells (DCs), and B cells to secondary lymphoid organs (SLOs) (3–7). Its two endogenous ligands, CC chemokine ligand 19 (CCL19) and 21 (CCL21), form chemotactic gradients within SLOs to guide CCR7-expressing cells (8–11). This homing mechanism is essential for bringing naïve T cells into close proximity with antigen-presenting DCs within the T cell zones of SLOs, a prerequisite for T cell priming and the initiation of antigen-specific immunity (12). Given its pivotal role, the CCR7 axis is a critical therapeutic target for a broad spectrum of human pathologies (13–15). Hyperactive CCR7 signaling promotes cancer metastasis and autoimmune disorders, whereas insufficient activity impairs immune surveillance and vaccine efficacy, highlighting the need for precisely tuned modulators.

However, effectively targeting this axis is complicated by the phenomenon of biased agonism, for which the CCR7 system is a paradigmatic example in the chemokine family (16). Although both ligands engage the same receptor, they induce strikingly different downstream signaling profiles and cellular responses (SI Appendix, Fig. S1B) (17–20). While sharing a structurally conserved core chemokine domain, CCL21 possesses a unique, highly basic C-terminal tail absent in CCL19 (SI Appendix, Fig. S1C). This tail facilitates binding to glycosaminoglycans (GAGs) on cell surfaces and in the extracellular matrix, which is crucial for establishing stable chemotactic gradients and facilitating haptotaxis (20–22). Beyond GAG interactions, their in vivo profiles are shaped by atypical chemokine receptors: CCRL2 selectively sequesters CCL19, whereas ACKR4 scavenges both (23–25). Nevertheless, the fundamental functional divergence is generally thought to be primarily driven by their biased agonism directly at CCR7.

Functionally, CCL19 potently induces G protein activation and β-arrestin recruitment. This process is mediated by receptor phosphorylation via GPCR kinases (GRKs), specifically GRK3 and GRK6, leading to potent but transient G protein signaling characterized by rapid receptor desensitization and internalization (17–19). In contrast, CCL21 activates G protein signaling with a potency initially reported to be comparable to, or approximately 10-fold lower than, that of CCL19, but it inefficiently recruits β-arrestin following phosphorylation solely by GRK6, resulting in minimal internalization (17–20). This differential regulation is physiologically crucial. The acute nature of CCL19 signaling facilitates localized cellular activation and precise positioning within SLOs, whereas the sustained signaling induced by CCL21, combined with GAG-regulated localization, establishes the stable, long-range chemotactic gradients required for guiding cell migration over distances (22, 26–28).

Despite its profound physiological importance, the mechanism by which these two distinct chemokines elicit biased signaling through CCR7 has been a long-standing enigma (29). Understanding the molecular mechanisms underlying this bias requires elucidating how ligand binding translates into distinct receptor conformations that selectively engage G proteins versus GRKs and arrestins. While the crystal structure of inactive-state CCR7 bound to the small-molecule intracellular antagonist Cmp2105 provided valuable insights into inhibitor binding (15), the structural underpinnings of its activation by endogenous chemokines remain elusive.

Here, we report the cryo-electron microscopy (cryo-EM) structures of human CCR7 in complex with CCL19 and CCL21, each stabilized in an active conformation by the heterotrimeric inhibitory G protein (G_i_). These structures reveal distinct ligand binding modes and differential receptor conformations induced by the two chemokines. The structures illustrate a unique mechanism of receptor activation shared within the CCR7-like chemokine receptor subfamily. Concurrently, while the overall G_i_ binding mode is similar, subtle conformational differences in intracellular regions not directly involved in G_i_ binding, such as ICL1, suggest distinct downstream signaling profiles. Motivated by these distinctions, we performed molecular dynamics (MD) simulations to detail the dynamic basis for these structural differences after G protein dissociation. These simulations revealed substantial differences in receptor conformational equilibria, especially in helix 8 (H8), that might influence GRK recruitment. Importantly, structure-guided mutagenesis targeting this intracellular network selectively impaired arrestin recruitment, supporting these computational findings and validating our hypothesis regarding the dynamic control of transducer selectivity. This study elucidates the structural basis for G protein activation by two distinct chemokines, providing fundamental insights into the mechanism of biased agonism at CCR7 and offering a structural template for the rational design of pathway-selective agonists.

## Results

### Preparation and overall architecture of the active CCR7-G_i_ complexes

To elucidate the structural basis of CCR7 signaling, we utilized a robust strategy to assemble stable, active-state complexes for cryo-EM (SI Appendix, Fig. S2 and Methods). We employed a temporally controlled expression system in tetracycline-inducible Expi293F (Expi293Fi) cells, expressing an engineered G_i_ heterotrimer prior to human CCR7 to maximize receptor stability and complex formation efficiency (SI Appendix, Fig. S2A). The G_i_ complex incorporated a single-chain Gγ-Gα_i_ fusion (30) with dominant negative Gα_i_ mutations (31) for enhanced stability. Additionally, the receptor and G protein were non-covalently tethered using the NanoBiT system (LgBiT fused to CCR7 and HiBiT fused to Gβ) (32, 33). The ligand-free CCR7-G_i_ complex, further stabilized by the fiducial marker scFv16 (34), was purified and the final signaling complexes were formed by adding separately purified full-length CCL19 or CCL21, yielding viable samples for cryo-EM analysis (SI Appendix, Fig. S2B-F).

Single-particle cryo-EM analysis of the samples yielded high-resolution maps with nominal resolutions of 3.0 Å for the CCL19-CCR7-G_i_-scFv16 complex and 3.2 Å for the CCL21-CCR7-G_i_-scFv16 complex (Fig. 1A, and SI Appendix, Table S1 and Fig. S3-S6). During 3D classification, we observed structural heterogeneity related to post-translational modifications (glycosylation at N36 and N292 of CCR7) and weak densities suggesting partial ordering of the proximal CCL21 C-terminal tail. However, as these classes represented minor populations and exhibited lower local resolution at the core interface, we focused on the most homogeneous particle subsets to accurately define the primary ligand binding modes.

**Fig. 1.**
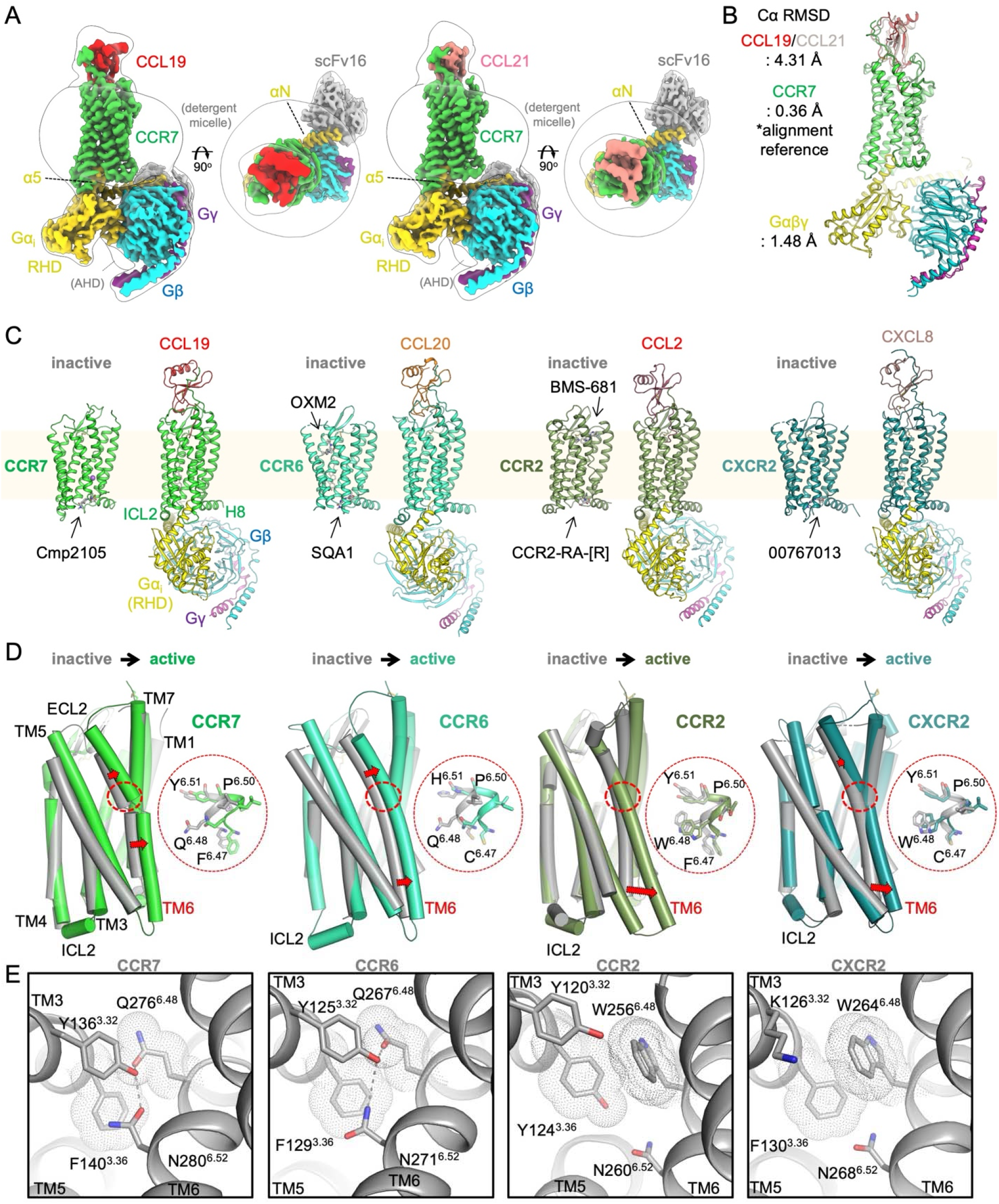
Structures of CCR7-G_i_ complexes and global conformational changes upon activation. (A) Cryo-EM maps of the CCL19-CCR7-G_i_-scFv16 (left) and CCL21-CCR7-G_i_-scFv16 (right) complexes. Structures are shown in side views and top views. Color coding: CCL19 (red), CCL21 (orange), CCR7 (green), Gα_i_ (yellow; RHD: Ras Homology Domain, AHD: Alpha Helical Domain), Gβ (cyan), Gγ (purple), scFv16 (gray). (B) Superposition of the ribbon models of the CCL19-bound and CCL21-bound complexes. Alignment was performed based on CCR7. Cα root-mean-square deviations (RMSD) are indicated for the chemokines, CCR7, and Gαβγ. (C) Comparison of active and inactive state structures of CCR7 and related chemokine receptors. Overall structures showing G protein coupling or antagonist/inhibitor binding. Structures include inactive CCR7 bound to Cmp2105 (PDB: 6QZH); active CCR7 bound to CCL19 (this study); inactive CCR6 bound to OXM2 and SQA1 (PDB: 9D3E); active CCR6 bound to CCL20 (PDB: 6WWZ); inactive CCR2 bound to BMS-681 and CCR2-RA-[R] (PDB: 5T1A); active CCR2 bound to CCL2 (PDB: 7XA3); inactive CXCR2 bound to 00767013 (PDB: 6LFL); active CXCR2 bound to CXCL8 (PDB: 6LFO). (D) Superposition of the inactive and active structures of chemokine receptors. Inactive states are shown in gray, and active states are separately colored. Red arrows indicate the outward movement of TM6. The region corresponding to the conserved CWxP motif (FQxP in CCR7 and CCR6) is indicated by red dashed circles and shown on the right. (E) Structural comparison of the inactive-state chemokine receptors viewed from the extracellular side. Key residues around position 6.48 are shown as sticks, and the F/Y^3.36^ and Q/W^6.48^ sidechains are displayed with dot surfaces to illustrate their steric bulk. Hydrogen-bonding networks are indicated by dashed lines.

The overall architecture of the complexes resembles previously reported GPCR-G protein structures, characterized by the canonical insertion of the Gα_i_-α5 helix into the receptor’s cytoplasmic core (Fig. 1A-C). The seven-transmembrane (7TM) conformations of CCR7 in both structures are essentially identical, with a Cα root-mean-square deviation (RMSD) of 0.36 Å. They also exhibit a highly similar docking mode for the G_i_ heterotrimer (Cα RMSD of 1.48 Å). Strikingly, however, the chemokines themselves adopt distinct orientations, reflected in a substantial Cα RMSD of 4.31 Å when aligning the complexes based on the receptor (Fig. 1B).

### A lateral TM6 movement during CCR7 activation

To understand the conformational changes associated with CCR7 activation, we compared the CCL19-bound active-state structure with the inactive CCR7 structure bound to the intracellular antagonist Cmp2105 (15) to elucidate the general mode of activation in comparison with other chemokine receptor systems (Fig. 1C). Both CCL19 and CCL21 induce a similar global conformational rearrangement in CCR7, creating a cytoplasmic cavity to accommodate the G protein.

The most prominent feature of this transition is the outward movement of transmembrane helix 6 (TM6). Unlike the canonical hinge-like pivot observed in most class A GPCRs, including CCR2 and CXCR2 (35, 36), CCR7 exhibits a predominantly parallel outward shift of TM6 by 3-4 Å (Fig. 1D). Specifically, the entire TM6 helix, from its extracellular end to its intracellular tip, moves laterally as a semi-rigid unit away from the receptor’s central axis. This activation mode is similar to that observed in its paralog, CCR6 (34% sequence identity), which also shows a ∼5 Å global shift (37).

These distinct activation modes correlate with the presence or absence of the critical W^6.48^ residue within the CWxP motif (superscripts denote Ballesteros-Weinstein numbering) (Fig. 1D). In most class A GPCRs, W^6.48^ acts as a rotamer “toggle switch,” initiating conformational changes that displace the intracellular half of TM6 (36, 38). In CCR7 and CCR6, this tryptophan is replaced by glutamine (Q^6.48^). We propose that this subclass-specific activation pathway is structurally encoded in the inactive state (Fig. 1E). In CCR7 and CCR6, the less bulky Q^6.48^ allows the extracellular portion of TM6 to pack closer to the receptor’s central axis, closely approaching the bulky F^3.36^ on TM3. Furthermore, a hydrogen-bond network centered on Y^3.32^ stabilizes the extracellular ends of TM3 and TM6 in close proximity. Upon activation, the insertion of the bulky chemokine core acts as a wedge, forcing the entire TM6 laterally outward while the extracellular end of TM5 concurrently closes inward to clasp the ligand. In contrast, in canonical receptors like CCR2 and CXCR2, the bulky W^6.48^ restricts TM6 from closely approaching F/Y^3.36^ due to steric hindrance, enforcing a wider extracellular gap between TM3 and TM6. Because the extracellular segment of TM6 is already pre-opened in the inactive state, chemokine binding induces minimal outward movement at this site. Instead, activation primarily triggers the microswitch-driven opening of the intracellular half, producing the classic pivot movement.

### Distinct binding modes of CCL19 and CCL21 dictated by CRS3 interactions

The most pronounced differences between the two active complexes are found at the extracellular ligand-recognition interface. Strikingly, the core domain of CCL21 is globally rotated by 17.6° relative to CCL19 when viewed from the side of the receptor (Fig. 2A). This rotation results in significantly altered interaction interfaces, with a larger buried surface area for CCL21 (1,640 Å^2^) compared to CCL19 (1,380 Å^2^).

**Fig. 2.**
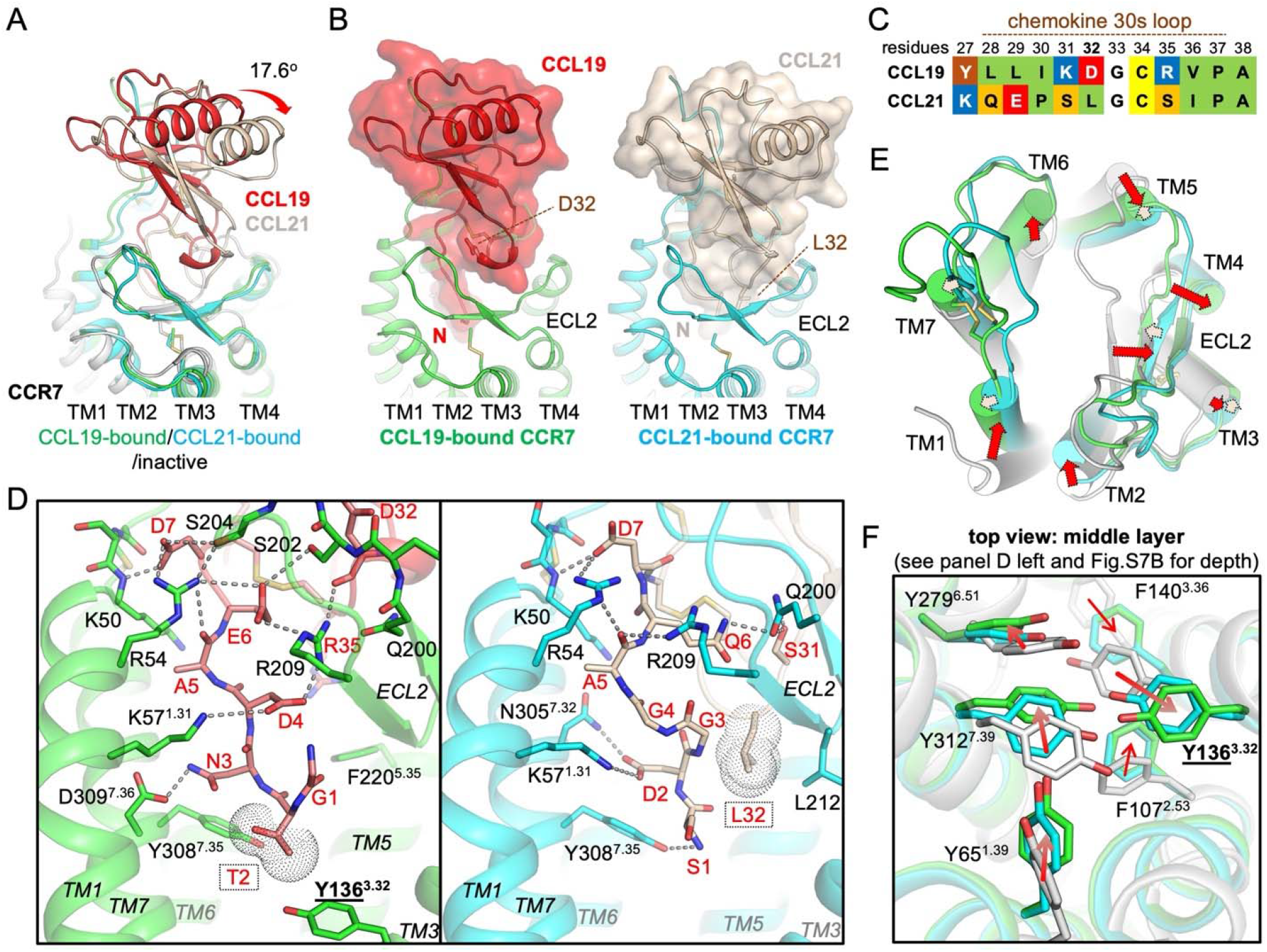
Distinct ligand binding modes of CCL19 and CCL21 at CCR7. (A) Superposition of the CCL19-CCR7 (red/green) and CCL21-CCR7 (orange/cyan) complexes with inactive CCR7 (gray), highlighting the 17.6° rotation of CCL21 relative to CCL19. (B) Divergent interactions at the CRS3 interface. CCL19-CCR7 (left) and CCL21-CCR7 (right) are shown side-by-side to compare the extracellular structures especially the interaction between the chemokine 30s loop and CCR7-ECL2 at CRS3. The central residues in the 30s loop (D32 for CCL19 and L32 for CCL21) are indicated as markers. (C) A sequence alignment of the 30s loops of CCL19 and CCL21. (D) Conserved and distinct interactions at the CRS2/3 interfaces. Close-up views of the chemokine N-terminus penetration into the orthosteric binding pocket for CCL19 (left) and CCL21 (right). Key interacting residues are shown as sticks, and polar interactions are indicated by dashed lines. (E) Ligand-specific conformational changes in the extracellular TMs and ECL2. Superposition of the CCL19-bound and CCL21-bound CCR7 structures with the inactive CCR7 structure, highlighting the conformational differences induced by the distinct ligand binding modes. Red arrows indicate conformational shifts from the inactive state to the CCL21-bound state, and peach arrows indicate the further shifts observed in the CCL19-bound state. (F) Rearrangement of the aromatic cluster in the middle layer of the TM bundle. Close-up view of the superimposed structures showing the distinct conformations of Y65^1.39^, F107^2.53^, Y136^3.32^, Y279^6.51^, Y308^7.35^, and Y312^7.39^ in the CCL19-bound and CCL21-bound states compared to the inactive CCR7 (PDB: 6QZH). See also Fig. S7.

Chemokine recognition generally follows an expanded “two-site” model (39), involving interactions across several chemokine recognition sites (CRS): CRS1 (receptor N-terminus/chemokine core), CRS1.5 (receptor N-loop and ECL3/chemokine N-loop), CRS2 (receptor 7TM pocket/chemokine N-terminus), and CRS3 (receptor ECL2/chemokine 30s loop) (SI Appendix, Fig. S7A). Our structures reveal that the dramatic difference in orientation stems primarily from divergent interactions at CRS3 (Fig. 2B).

Despite overall sequence conservation (32% identity, 68% similarity in the core region; SI Appendix, Fig. S1C), the 30s loops of CCL19 and CCL21 exhibit distinct sequences and conformations (Fig. 2B,C). The CCL21 loop (^28^QEPS**L**GCSIP^37^) is notably less bulky and more hydrophobic at the center than the corresponding loop in CCL19 (^28^LLIK**D**GCRVP^37^), featuring L32 instead of D32 (Fig. 2C). Consequently, the 30s loop of CCL19 adopts a curled conformation and rests atop CCR7-ECL2. In contrast, the more compact 30s loop of CCL21 penetrates deeper into the extracellular pocket. Crucially, CCL21-L32 inserts like a wedge between the chemokine N-terminus and CCR7-L212^ECL2^ (Fig. 2B,D). This deep insertion stabilizes a specific conformation of ECL2 and contributes to the overall rotation of CCL21.

In contrast to the divergent CRS3 interactions, the engagements at CRS1 and CRS2 are relatively conserved. At CRS1, CCR7-F44 forms the core of the interactions with the chemokine bodies (SI Appendix, Fig. S7B). At CRS2, the N-terminal loops of both chemokines penetrate the “minor orthosteric binding pocket” of CCR7 surrounded by TM1, TM2, TM3, TM6, and TM7, forming extensive polar and hydrophobic interactions (Fig. 2D) (40). The N-terminus of CCL19 penetrates slightly deeper into the pocket than that of CCL21. Specifically, the sidechain of CCL19-T2 engages Y136^3.32^ at TM3, inducing a subtle shift of this residue. The observed CRS2 interaction networks generally correlate with reported receptor mutagenesis data (41). For example, CCR7 residues K50^N-term^ and R54^N-term^ are located at the tip of TM1 and the center of the molecular interface, forming critical contacts with the N-terminal peptides of both chemokines (Fig. 2D). This explains why the CCR7-K50A^N-term^ and CCR7-R54A^N-term^ mutations dramatically impair both CCL19- and CCL21-induced signaling. Conversely, the CCR7-R209A^ECL2^ mutation selectively reduces CCL19-induced signaling. This selectivity is structurally rationalized by the more extensive polar contacts of R209^ECL2^ with the N-terminus/core of CCL19, via the D4/E6 sidechains and D32 backbone, compared to CCL21 (Fig. 2D), highlighting its specific role in CCL19 recognition.

These distinct binding modes result in ligand-specific conformational shifts in the extracellular portions of TM1, TM3, TM5, TM7, and ECL2 (Fig. 2E and SI Appendix, Fig. S7C). Furthermore, these differences propagate to an aromatic cluster located in the middle layer of the minor pocket, just beneath the chemokine N-termini (Fig. 2F and SI Appendix, Fig. S7B). Residues Y65^1.39^, Y136^3.32^, Y279^6.51^, and Y312^7.39^ exhibit distinct conformations in the inactive, CCL21-bound, and CCL19-bound states, with the CCL19-bound state exhibiting consistently more extensive changes. Among these residues, the Y312^7.39^E mutation is known to confer a high degree of constitutive activity to CCR7 while abolishing chemokine sensitivities, whereas Y312^7.39^A only causes high basal activities, underscoring the importance of this residue for the inactive-to-active transition (20). The correlated rearrangements in the aromatic network likely contribute to the differential stabilization of active-state conformations and may underlie the observed differences in general signaling potency (CCL19 > CCL21) (20).

### Structural basis of CCR7 activation and core G protein interface

The binding of both chemokines induces canonical conformational changes associated with class A GPCR activation (38) (Fig. 3A), although with the unique features related to TM6 movement described earlier. The PIF-like motif (P^5.50^-M^3.40^-F^6.44^) undergoes reorganization, characterized by the movement of the M144^3.40^ sidechain, which induces inward and outward shifts of TM5 and TM6, respectively (Fig. 3A, bottom left). The immediately downstream hydrophobic lock collapses to further destabilize the inactive state (Fig. 3A, bottom center).

**Fig. 3.**
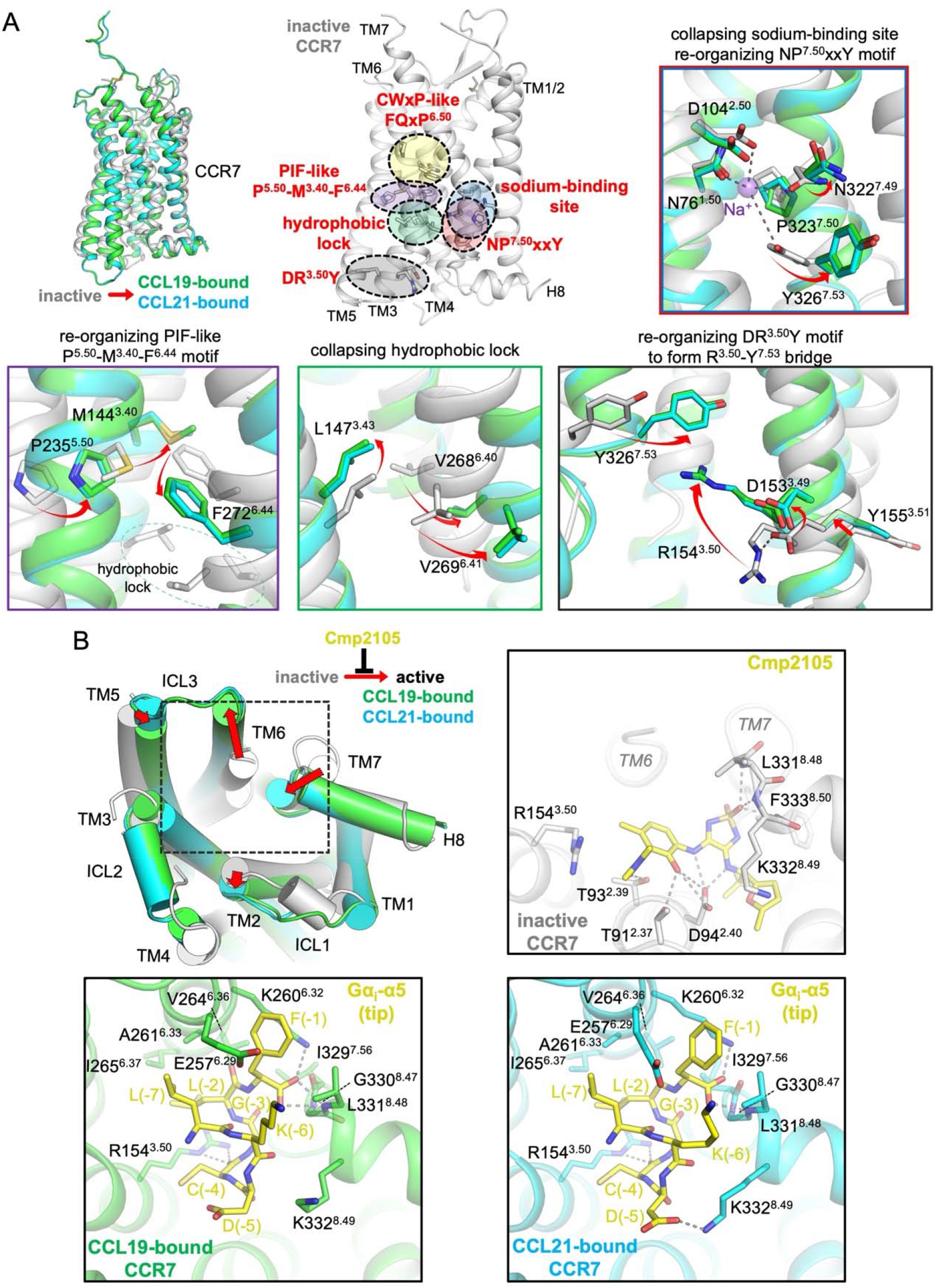
Structural basis of CCR7 activation. (A) Conserved GPCR activation motifs and conformational changes in CCR7. Top left: Superposition of inactive CCR7 (gray) and active CCR7 (CCL19-bound in green, CCL21-bound in cyan). Top center: Mapping of the key GPCR activation motifs highlighted on the inactive CCR7 structure. Other panels show close-up views of the reorganization of the motifs upon activation. Key residues are shown as sticks. Arrows indicate the direction of movement from the inactive to the active state. Top right: Collapse of the sodium-binding site and reorganization of the NP^7.50^xxY motif. Bottom left: Reorganization of the PIF-like P^5.50^-M^3.40^-F^6.44^ motif upstream of the hydrophobic lock. Bottom center: Collapse of the hydrophobic lock. Bottom right: Reorganization of the DR^3.50^Y motif and formation of a G protein-binding interface with the R^3.50^-Y^7.53^ bridge. (B) G protein coupling and comparison with the antagonist-bound state. Top left: Superposition of active CCR7 (CCL19-bound in green, CCL21-bound in cyan) and inactive CCR7 bound to Cmp2105 (gray). The inactive-to-active TM movements, which are generally conserved between the CCL19-bound and CCL21-bound states in complex with G_i_, are indicated by red arrows. Top right: Close-up view showing how Cmp2105 (yellow sticks) occupies the G protein binding site in the inactive CCR7. Cmp2105 also inhibits the inactive-to-active conformational transition. Bottom panels: Detailed interactions between the Gα_i_ α5 helix (yellow, C-terminal 7 residues) and the CCR7 intracellular cavity in the CCL19-bound (left) and CCL21-bound (right) complexes. Key residues involved in the interaction are shown as sticks, and polar contacts are indicated by dashed lines.

Concurrently, the sodium-binding site, which is stabilized by sodium ion coordination with N76^1.50^, D104^2.50^, N322^7.49^, and Y326^7.53^ in the inactive state, collapses upon activation, accompanied by the kinking of the NP^7.50^xxY motif at TM7 and the flipping of the Y326^7.53^ sidechain (Fig. 3A, top right). At the same time, the DR^3.50^Y motif reorganizes, allowing R154^3.50^ to form a bridge with Y326^7.53^, thereby stabilizing the active conformation for G protein coupling (Fig. 3A, bottom right).

Comparing these active states with the Cmp2105-bound inactive structure highlights the structural rearrangements required for G protein engagement (Fig. 3B, top left). The outward movement of TM6 and the concomitant shift of TM5 create the binding pocket for the Gα_i_ subunit, prompting significant reorganization of the intracellular loops. Notably, ICL2 transitions from a disordered loop in the inactive state to a stable, short α-helix upon activation, which is critical for engaging the Gα subunit. Similarly, ICL3 is stabilized following TM6 displacement to further form the G protein interface. Cmp2105 stabilizes the inactive state by preventing this conformational shift and sterically occluding the binding site of the Gα_i_-α5 helix (Fig. 3B, top right).

The primary interface between CCR7 and Gα_i_ involves the insertion of the C-terminal α5-helix of Gα_i_ deep into the cytoplasmic cavity of the receptor (Fig. 1A-C and 3B). This highly conserved interface is stabilized by extensive interactions in both the CCL19- and CCL21-bound complexes (Fig. 3B, bottom panels). The Gα_i_-F(−1) [F354] sidechain is sandwiched between E257^6.29^ and K260^6.32^, positioning its C-terminal carboxyl group for polar interactions with the backbone amide at the TM7-H8 turn and the K260^6.32^ sidechain. The L(−2) [L353] sidechain inserts into a hydrophobic pocket formed by TM6 residues and R154^3.50^ at TM3, capped by L(−7) [L348]. Additionally, the R154^3.50^ sidechain forms polar interactions with the C(−4) [C351] backbone carbonyl. Despite minor variations at the periphery of the core interface, the overall interaction patterns remain highly conserved.

### Subtle differences at the intracellular loops suggest distinct signaling outcomes

Despite the overall similarity in G protein coupling, a detailed comparison reveals subtle but notable differences in the intracellular loops between the CCL19- and CCL21-bound complexes (Fig. 4A,B).

**Fig. 4.**
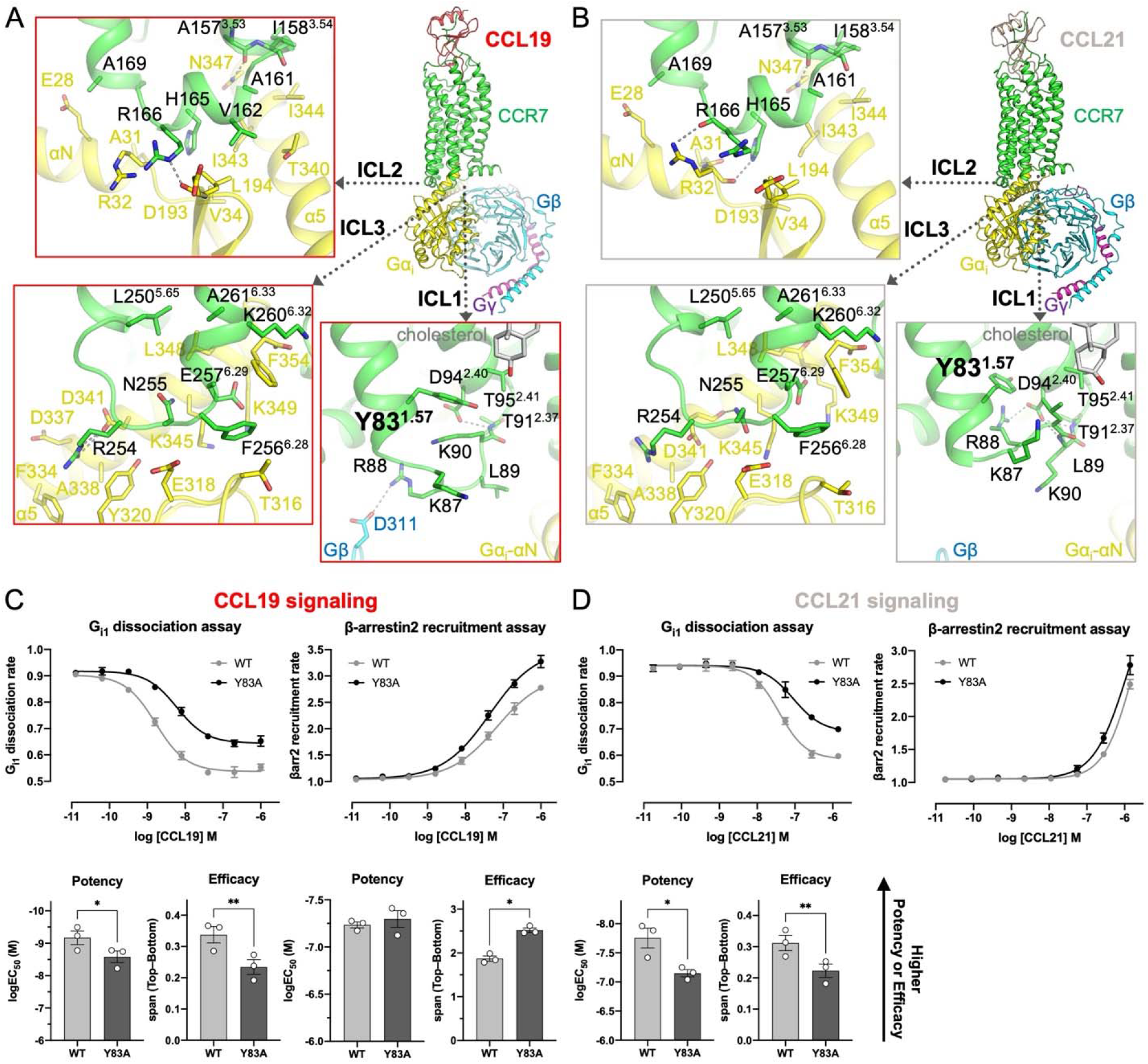
Conserved G_i_ interface with distinct Y83 rotameric states. The G_i_ binding interface of the (A) CCL19-CCR7-G_i_ and (B) CCL21-CCR7-G_i_ complexes. Overall views are displayed on top right, where zoomed-in boxes highlight the interfaces centering around ICL2 (top left), ICL3 (bottom left), and ICL1 (bottom right). The contacting residues within 4 Å distance are shown as sticks, and polar interactions are shown by dashed lines. Cholesterol molecules near ICL1 are shown as gray sticks. (C,D) Effects of the Y83A mutation on G_i1_ and β-arrestin2 signaling induced by CCL19 and CCL21. G_i1_ dissociation and β-arrestin2 recruitment in response to (C) CCL19 or (D) CCL21 at WT and Y83A CCR7. Representative concentration-response curves are shown in the top panels, while potency (logEC_50_) and efficacy (span; Top-Bottom) are presented as bar graphs in the bottom panels. Because a clear plateau was not achieved for β-arrestin2 recruitment in response to CCL21, logEC_50_ and span values could not be reliably estimated and are therefore not determined. Concentration-response curves represent a single experiment performed in triplicate and are shown as mean ± standard deviation (SD). Summary data represent three independent experiments and are shown as mean ± standard error of the mean (SEM). Statistical comparisons between WT and Y83A were performed using paired two-tailed t-tests (* *P* < 0.05; ** *P* < 0.01). βarr2, β-arrestin2.

The conformations of ICL2 and ICL3, which form the major part of the G_i_ interface, are essentially identical in both active structures. In the ICL2 region, the interactions between CCR7 residues (e.g., H165, R166) and the Gα_i_ αN-β1 hinge region (e.g., R32, D193) are conserved, exhibiting minor variations (Fig. 4A,B, top left panels). Similarly, the core interface mediated by ICL3, TM5, and TM6 is highly conserved (Fig. 4A,B, bottom left panels).

In contrast, ICL1, which is less directly involved in the primary G_i_ interface, adopts distinct loop conformations in the two complexes (Fig. 4A,B, bottom right panels). This divergence appears to originate from differences in the side-chain conformation of Y83^1.57^, located at the cytoplasmic end of TM1. The distinct binding modes of CCL19 and CCL21 induce specific rearrangements in the extracellular TMs (Fig. 2D) and the central aromatic network (Fig. 2F). This network appears to function as an allosteric link, propagating these differences through the helical bundle (particularly TM1) to induce the distinct rotameric states of Y83^1.57^ in the CCL19-versus CCL21-bound structures. These states influence the length of TM1 and the positioning of adjacent residues, propagating to the overall conformation of the ICL1 loop.

To investigate the functional relevance of the observed structural differences, we first evaluated the signaling profiles of N-terminally FLAG-tagged wild-type (WT) CCR7 using NanoBiT-based G_i1_ dissociation and β-arrestin2 membrane recruitment assays (SI Appendix, Fig. S8A). Given the distinct rotameric states of Y83^1.57^ observed between the two active structures (Fig. 4A,B), we next examined the impact of the Y83^1.57^A mutation on this signaling balance (Fig. 4C, D). In response to CCL19, the Y83^1.57^A mutant exhibited reduced potency and efficacy for G_i1_ activation, but maintained potency and elevated efficacy for β-arrestin2 recruitment, despite comparable cell surface expression (SI Appendix, Fig. S8B). Consequently, the Y83^1.57^A mutation shifts the signaling preference toward the β-arrestin2 pathway, suggesting that the Y83^1.57^-mediated conformational transition is critical for transducer selection.

Thus, while the core G_i_-binding interfaces are highly conserved, our functional data demonstrate that structural variations around the Y83^1.57^ microswitch impact signaling preference. While ICL1 is expected to be flexible due to its lack of direct G protein engagement, the observation of distinct loop conformations even within the static cryo-EM structures highlights the need to investigate the dynamic behavior of the receptor (SI Appendix, Fig. S5C,D and S6). Although ICL1 itself may not be the primary GRK binding site, we hypothesize that the distinct ligand binding modes, relayed via Y83^1.57^, prime the receptor core and intracellular loops to adopt distinct conformational ensembles in the absence of the G protein (42, 43).

### Molecular dynamics simulations to probe the basis of biased signaling

To investigate how the distinct ligand binding modes and the subtle intracellular differences might influence receptor dynamics relevant to biased signaling, we performed six independent 2.3 μs all-atom MD simulations for each complex (Fig. 5A and SI Appendix, Fig. S9-S12). We used the experimental CCL19- and CCL21-bound structures as starting points, removing the G protein and scFv16 and embedding them in a lipid bilayer to mimic the receptor states primed for subsequent signaling events, such as GRK recruitment (SI Appendix, Fig. S9A). During the simulations, the chemokines exhibited high flexibility, consistent with the cryo-EM densities (SI Appendix, Fig. S3, S4, and S9B,C). Conversely, the receptor 7TM bundles slightly deviated from their initial conformations but converged to similar overall states (SI Appendix, Fig. S9B,C and S10).

**Fig. 5.**
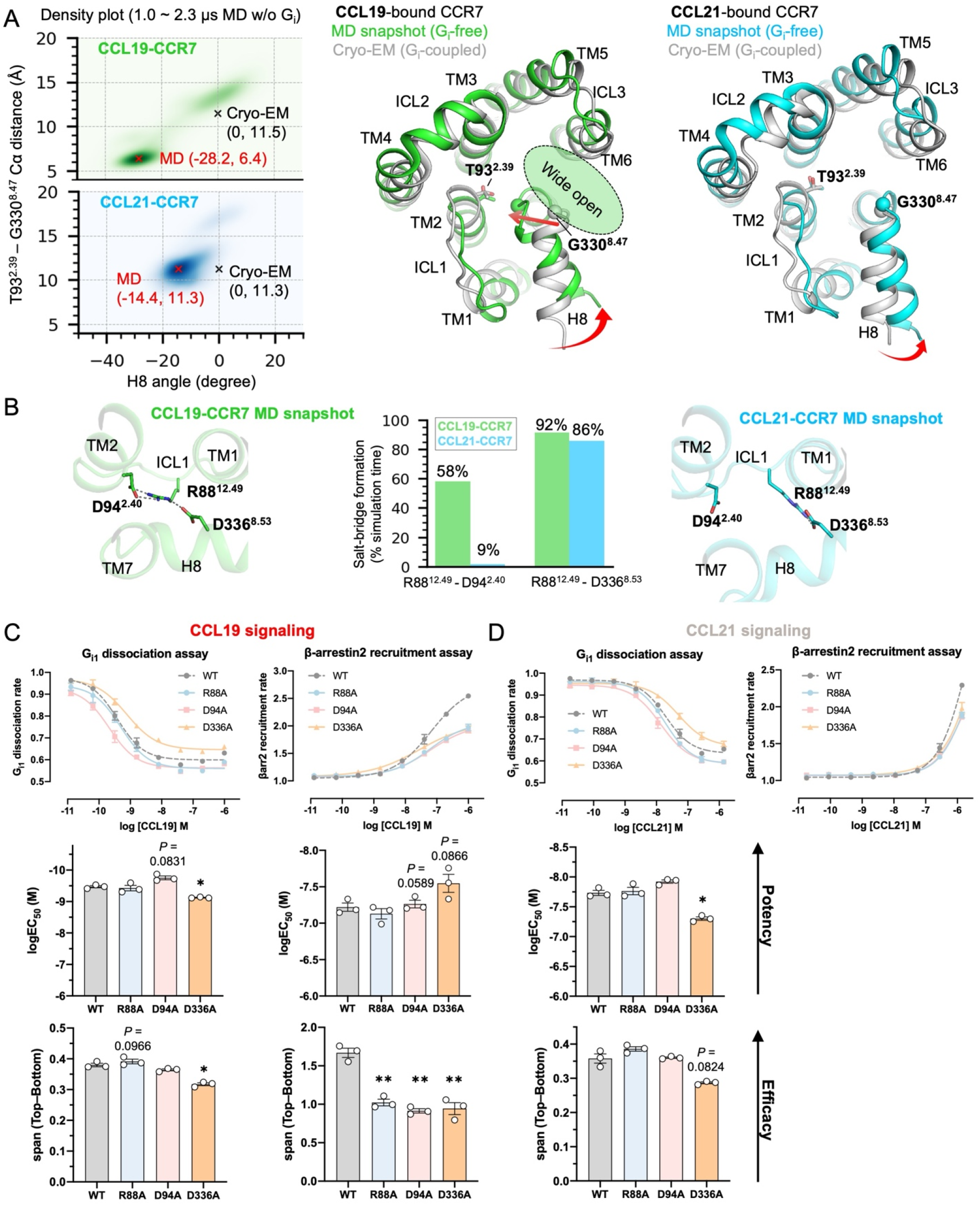
Distinct intracellular dynamics revealed by MD simulations and functional validation. (A) Comparison of the H8 orientation after the G_i_ free MD simulations between CCL19-bound (green) and CCL21-bound (cyan) CCR7. Left: Probability density plot of the H8 angle versus the T93^2.39^-G330^8.47^ distance. Data were calculated from six independent 2.3 μs G_i_-free MD simulations, excluding the first 1 μs of each run as a relaxation period. See Fig. S11C for the definition of the H8 angle. The angles and the distances from the cryo-EM models and the MD density peak are indicated by black and red crosses (‘x’) with corresponding text, respectively. Right: Superposition of the G_i_ free MD snapshot and the G_i_-coupled cryo-EM structure. The residues used for the distance measurement in the density plot, T93^2.39^ and G330^8.47^, are shown as sticks and spheres, respectively. The larger degree of the H8 rotation and the consequent insertion of the TM7-H8 loop open the gap between H8 and TM6 in the CCL19-bound CCR7 compared to the CCL21-bound CCR7. (B) Structural basis of the distinct H8 orientation between CCL19-bound CCR7 and CCL21-bound CCR7 in the G_i_ free MD simulations. The closer positioning of H8 towards ICL1 and TM2 in CCL19-bound CCR7 is stabilized by the salt-bridge network of R88^12.49^ to D94^2.40^ and D336^8.53^. The frequency of the salt-bridge formation between R88^12.49^ (ICL1) - D94^2.40^ (TM2) and between R88^12.49^ (ICL1) - D336^8.53^ (H8) during the MD simulations is shown (middle, the salt-bridge distance threshold of 4 Å). A close-up view of H8, TM1, ICL1, and TM2 in the MD snapshots of CCL19-bound CCR7 (left) and CCL21-bound CCR7 (right). The residues of interest are shown as sticks, and salt bridges are shown in dashed lines. (C,D) Effects of the salt bridge mutations on G_i1_ and β-arrestin2 signaling induced by CCL19 and CCL21. G_i1_ dissociation and β-arrestin2 recruitment in response to (C) CCL19 or (D) CCL21 at WT, R88A, D94A, and D336A CCR7. Representative concentration-response curves are shown in the top panels, while potency (logEC_50_) and efficacy (span; Top-Bottom) are presented as bar graphs in the bottom panels. Because a clear plateau was not achieved for β-arrestin2 recruitment in response to CCL21, logEC_50_ and span values could not be reliably estimated and are therefore not determined. Concentration-response curves represent a single experiment performed in triplicate and are shown as mean ± SD. Summary data represent three independent experiments and are shown as mean ± SEM. Statistical analysis was performed using one-way ANOVA followed by Dunnett’s multiple comparisons test versus WT (* *P* < 0.05; ** *P* < 0.01). Exact *P* values are shown when 0.05 < *P* < 0.1.

The most notable finding was the differential dynamics of H8 in CCR7 (Fig. 5A). In the CCL19-bound state, the TM7-H8 loop is highly dynamic, repositioning toward the receptor central axis and widening the gap between TM6 and TM7 (Fig. 5A). This shift is characterized by the approach of the TM7-H8 loop toward TM2 and a pronounced H8 rotation (SI Appendix, Fig. S11). Specifically, the Cα distance between T93^2.39^ and G330^8.47^ narrowed substantially from 11.5 Å to 6.4 Å, accompanied by an H8 rotation to a new stable state at -28.2° (relative to the experimental structure, defined as 0°). In contrast, H8 in the CCL21-bound state remained stable near the cryo-EM conformation, with the corresponding Cα distance remaining at 11.3 Å and a less pronounced rotation of -14.4°.

These differences trace back to the formation of two distinct salt bridges: a highly stable one directly linking R88^12.49^ (ICL1) to D336^8.53^ (H8) (present in 92% of simulation time), and a second, dynamic one between R88^12.49^ (ICL1) and D94^2.40^ (TM2) (present in 58% of simulation time) (Fig. 5B and SI Appendix, Fig. S12B). In sharp contrast, in the CCL21-bound state, while the R88^12.49^-D336^8.53^ bridge remained largely stable (86% formation), the crucial R88^12.49^-D94^2.40^ bridge, which brings H8 toward TM2, was almost entirely absent (9% formation).

This dynamic behavior in the CCL19-bound state also correlates with the different chi1 (χ1) angles of Y83^1.57^ at the TM1 terminus (180° for CCL19 and -86° for CCL21), observed in the cryo-EM structures, which remained stable in their respective states throughout the simulations (SI Appendix, Fig. S12C). This orientation of Y83^1.57^ appeared to influence the ICL1 dynamics and allosteric coupling between ICL1 and H8 (Fig. 5A,B and SI Appendix, Fig. S12D). In the CCL19-bound state, the outward-facing conformation of Y83^1.57^ accommodates F333^8.50^ from H8 into a pocket near TM2, whereas the inward-facing Y83^1.57^ in the CCL21-bound state sterically blocks this insertion (SI Appendix, Fig. S12A).

Consequently, the stable conformational ensemble induced by CCL21, which lacks deep H8 insertion and resembles the initial G_i_-bound conformation, appears suboptimal for GRK interaction. This “locked” state facilitates G protein-biased signaling. Conversely, the conformational flexibility induced by CCL19 allows the receptor to sample multiple states, including a major “open” population characterized by deeper TM7-H8 loop insertion, alongside a population resembling the starting Gi-favorable state. This dynamic equilibrium likely represents the structural basis for CCL19’s balanced activation profile, enabling efficient engagement of both effectors.

### Functional assessment of the dynamic states dictating effector selection

To evaluate this MD-derived dynamic model, we assessed both the G_i1_ and β-arrestin signaling profiles of three structure-guided CCR7 mutants: R88^12.49^A, D94^2.40^A, and D336^8.53^A (Fig. 5C, D). All mutants exhibited cell surface expression levels statistically comparable to WT (SI Appendix, Fig. S8B). Notably, all three mutations significantly and uniformly reduced the maximal efficacy of CCL19-induced β-arrestin2 recruitment (Fig. 5C), highlighting the strict structural requirements for the arrestin-competent state. However, their divergent effects on G_i1_ signaling perfectly encapsulate our dynamic switching model. Similar overall trends were also observed upon CCL21 stimulation (Fig. 5D).

The MD simulations indicated that D336^8.53^ serves as a stable resting point for R88^12.49^ in both CCL19- and CCL21-bound states (Fig. 5B). Consistent with the role of D336^8.53^ as a fundamental structural anchor for the receptor’s architecture, the D336^8.53^A mutation significantly reduced overall receptor function, demonstrating reduced potency (an increase in the EC_50_) and a reduction in efficacy for G_i1_ activation, alongside the aforementioned reduction in β-arrestin2 recruitment efficacy (Fig. 5C).

Conversely, adopting the GRK-favorable “open” state involves R88^12.49^ dynamically engaging with D94^2.40^ and pulling D336^8.53^ on H8 inward. This transition is highly populated in the balanced CCL19 complex but nearly absent in the G protein-biased CCL21 complex (Fig. 5B). Disrupting this “open-state” tether with the D94^2.40^A mutation resulted in the expected reduction in β-arrestin2 efficacy (Fig. 5C). By limiting the receptor’s ability to sample this “open” conformation, the D94^2.40^A mutation appears to favor the “locked” G_i_-selective state, thereby maintaining robust WT-level G_i1_ activation despite the targeted loss of β-arrestin2 efficacy.

The role of R88^12.49^ in this dynamic mechanism is further supported by the R88^12.49^A mutant. Removing this central basic residue likely impairs the receptor’s transition into the D94^2.40^-tethered “open” state, which is consistent with the significant reduction in β-arrestin2 efficacy (Fig. 5C). However, its absence does not disrupt the fundamental G_i_-binding pocket, allowing the receptor to adopt a G_i_-favorable conformation and sustain WT-like G_i1_ activation. Together, these functional data suggest that the dynamic R88^12.49^-mediated network contributes to transducer selection at CCR7, and that the transition required for efficient arrestin engagement depends on this intact basic anchor.

## Discussion

The precise orchestration of immune cell migration by the CCL19/CCL21-CCR7 axis relies on biased agonism for localized activation versus long-range chemotaxis (22, 26, 27, 44). Here, our structures suggest a molecular framework for this functional divergence. We reveal that ligand recognition is primarily dictated by interactions at the CRS3 interface (Fig. 2). The deep, hydrophobic engagement of the CCL21 30s loop contrasts sharply with the shallower binding of the CCL19 loop, illustrating how the sequence variations in flexible chemokine loops can alter the overall binding topology. Furthermore, our analysis highlights that CCR7 employs a non-canonical, rigid-body movement of TM6 during activation (Fig. 1), likely facilitated by the absence of the canonical W^6.48^ toggle switch, suggesting a specialized adaptation within this receptor subfamily for transducing signals from protein ligands.

The central paradox of CCR7 bias is that both ligands induce highly similar G_i_-coupled conformations (Fig. 4A,B) yet elicit vastly different downstream responses regarding desensitization (19). Our MD simulations suggest that the mechanism of bias lies not only in the initial G protein engagement, but also in the receptor dynamics that follow G protein dissociation, revealing that CCL19 induces a flexible ensemble capable of adopting a widely “open” H8 state, whereas CCL21 stabilizes a single, “locked” G_i_-bound-like state (Fig. 5A).

Structural comparisons with GPCR-GRK complexes, such as neurotensin receptor 1 (NTSR1) and Rhodopsin (Rho), underscore the importance of this CCL19-induced “open” state (Fig. 6). These structures reveal that the binding modes of the Gα C-terminal α5 helix and the GRK N-terminal α-helix (αN) are mutually exclusive (42, 43, 45, 46) (Fig. 6A, B). While the Gα-α5 helix requires deeper insertion towards the receptor core, the GRK-αN helix exhibits a shallower insertion and exits through the lateral gap between TM6 and the TM7-H8 loop. This suggests a general mechanism whereby efficient GRK recognition requires both a sufficiently wide lateral opening and a conformation that disfavors deep G protein re-insertion.

**Fig. 6.**
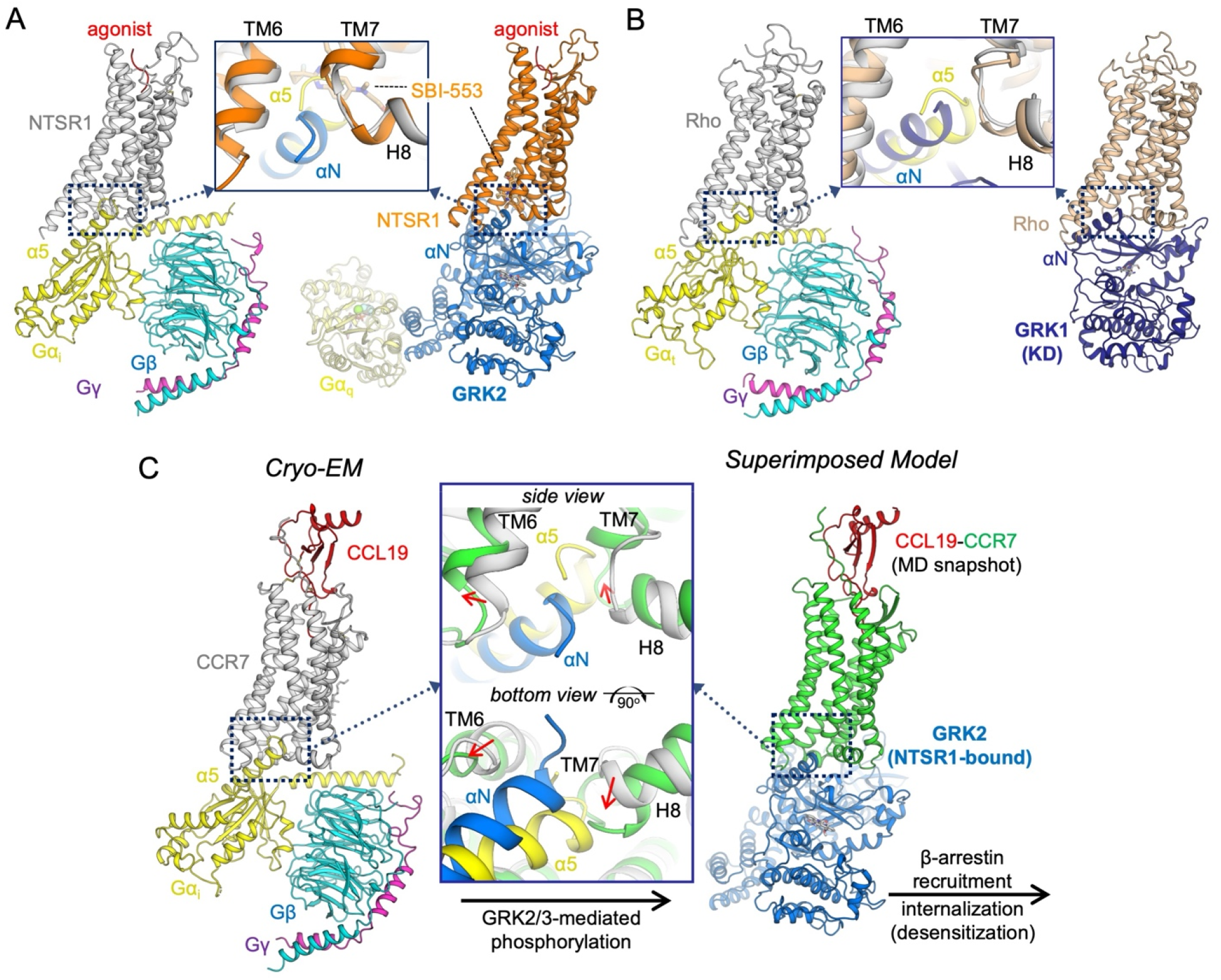
Comparison of G protein and GRK binding modes and a structural model for CCR7 biased agonism. (A) Cryo-EM structures of Neurotensin Receptor 1 (NTSR1) in complex with G_i_ (PDB: 6OS9, left) and GRK2 (PDB: 8JPB, right). The zoomed-in inset shows a superposition of the two complexes, aligned on the receptor (NTSR1), to directly compare the binding modes of the C-terminal α5 helix of the Gα subunit (yellow) and the N-terminal αN helix of the GRK (blue). (B) Cryo-EM structures of Rhodopsin (Rho) in complex with G_t_ (PDB: 6OY9, left) and GRK1 (PDB: 7MTA, right). The zoomed-in inset shows a superposition of the Rho-G_t_ and Rho-GRK1 complexes, aligned on the receptor (Rhodopsin), to compare the effector helices. Both (A) and (B) highlight that while both effectors engage the receptor’s cytoplasmic core, the GRK-αN helix exhibits a shallower insertion than the Gα-α5 helix (yellow) and follows a distinct trajectory, exiting through the interface formed by TM6, TM7, and H8. (C) Proposed structural model for the biased activation of GRK2/3 by CCR7. Cryo-EM structure of the CCL19-CCR7-G_i_ complex (this study, left) and a superimposed model illustrating the proposed mechanism of CCR7-GRK2 engagement (right). The model was generated by superimposing a representative “open” conformation of CCL19-CCR7 (green), derived from MD simulations (Fig. 5A), onto the NTSR1 receptor of the cryo-EM NTSR1-GRK2 complex (PDB 8JPB). The zoomed-in inset compares the G_i_-bound state (gray helices, Gα-α5 in yellow) with the modeled ‘open’ GRK-bound state (green helices, GRK-αN in blue). This illustrates that the “open” state, stabilized by CCL19-induced H8 dynamics after G protein dissociation, creates a wide gap between TM6 and H8, which appears sterically complementary to accommodate the GRK-αN helix. This simple model suggests that CCL19-induced dynamics selectively favor this “open” ensemble, facilitating efficient GRK2/3 recruitment and robust β-arrestin signaling.

In this context, the conformational flexibility of CCL19-bound CCR7 provides a compelling structural rationale for its balanced signaling. Its ability to dynamically adopt the “open” state, where the TM7-H8 loop inserts into the cytoplasmic cavity (Fig. 5A), simultaneously achieves two goals. It sterically hinders deep G protein re-engagement while creating a widened lateral gap favorable for GRK3 engagement (Fig. 6C). Furthermore, this inserted TM7-H8 loop may form a novel composite interaction surface for GRK recognition, analogous to how the GRK-biased small molecule SBI-553 acts on NTSR1 (43). This dynamic equilibrium permits efficient engagement of both effector types. Conversely, the “locked” state stabilized by CCL21 maintains a vacant deep pocket and a narrower lateral gap, rendering it competent for G protein coupling but deficient in the steric opening or interaction surface required for efficient GRK3 engagement. This structural dichotomy offers a mechanistic basis for CCL21’s G protein bias.

Our mutagenesis data provide functional support for this dynamic model (Fig. 4 and 5), wherein ligand-specific conformations are relayed via the Y83^1.57^ microswitch to govern the intracellular D94^2.40^-R88^12.49^-D336^8.53^ network. Disrupting specific nodes within this network predictably shifts signaling preference: the Y83^1.57^A mutation favors the arrestin pathway, whereas the D94^2.40^A and R88^12.49^A mutations restrict the receptor to a G_i_-favorable state.

The conceptual framework parallels recent findings in the µ-opioid receptor (µOR), where structural rearrangements narrow the intracellular core to favor arrestin binding over G protein coupling (47). Unlike the concurrent inward shift of TM7 and the TM7-H8 loop in µOR, CCR7 maintains a stable TM7 backbone while its TM7-H8 loop dynamically shifts inward. Pharmacologically, µOR exhibits a pronounced, switch-like shift in signaling bias, whereas CCR7 relies on subtle fine-tuning. Despite these differences, both share a fundamental structural principle: partial occupation of the receptor core cavity sterically hinders canonical G protein re-engagement while accommodating the arrestin pathway.

Physiologically, CCL19 drives rapid desensitization and internalization suited for acute immune tuning and tissue development (17, 19, 26, 44, 48–51). In contrast, the conformationally restricted ensemble of CCL21 disfavors efficient GRK3 interaction, resulting in sustained G protein signaling essential for long-distance immune cell guidance (17, 22, 28, 49). Moreover, the deep insertion of the CCL21 30s loop may contribute to a slower dissociation rate, further supporting sustained signaling, which, alongside GAG-mediated gradient formation (20–22), explains its role in long-range haptotaxis. In conclusion, this structural and dynamic framework reveals how subtle differences in ligand recognition translate into distinct physiological outcomes. Exploring chemokine analogs that mimic the CRS2/CRS3 engagement profile of CCL21 but at much higher affinity might lead to G protein-biased agonists. Such molecules could promote sustained cell trafficking as adjuvants for vaccines or cancer immunotherapy, providing a foundation for novel immunomodulators.

### Limitations of Study

While the current study provides significant insights into CCR7 activation and biased signaling mechanisms, several limitations remain. First, our structures do not directly visualize the conformations responsible for GRK or β-arrestin engagement; determining these complex structures will be crucial to fully elucidate the mechanisms of bias. Second, although we observed heterogeneity related to the CCL21 C-terminal tail and potential post-translational modifications, we could not resolve these features at high resolution, warranting further structural investigation. Finally, because of nonspecific activity at 3 µM, CCL21 concentrations in our functional assays were restricted to 1.39 µM. While the G_i1_ assay yielded reliable data within this range, the β-arrestin2 assay did not reach a plateau, precluding the estimation of EC_50_ and maximal responses (E_max_) and limiting our conclusions regarding CCL21-mediated β-arrestin2 signaling.

## Materials and Methods

### Mammalian cell culture for recombinant protein production

Expi293F inducible (Expi293Fi) cells (Thermo Fisher Scientific, a component of expression system kit Cat# A39251) were adopted and maintained in HE400AZ media (GMEP) supplemented with 10 µg/mL Blasticidin (InvivoGen) at a 20 mL culture scale. For protein expression, cells were expanded with HE400AZ media up to 200 mL and then HE200 media (GMEP) supplemented with GlutaMAX (Thermo Fisher Scientific) for larger volumes to reduce the total cost. Cells were maintained at a density between 0.3 × 10^6^/mL and 4 × 10^6^/mL and shaken at 37°C and 130 rpm with 8% (v/v) CO_2_. Transfections were performed at a cell density of approximately 3 × 10^6^/mL.

### Insect cell culture for recombinant protein production

For scFv16 expression from *Trichoplusia ni* (Hi5) ovarian cells (Expression Systems, Cat# 94-002), baculovirus was produced in *Spodoptera frugiperda* (Sf9) ovarian cells (ATCC CRL-1711) maintained in Sf-900 III medium (Thermo Fisher Scientific) with 10% (v/v) FBS (Sigma-Aldrich) and 2 mM L-Gln. Hi5 cells were maintained and expanded for protein expression in ESF 921 Insect Cell Culture Medium (Expression Systems). Insect cells were grown at 27°C with ambient CO_2_ and gentle agitation, and baculovirus infections for virus amplification and protein expression were performed at a cell density of approximately 2 × 10^6^/mL.

### Expression of the apo CCR7-G_i_ complex

The expression plasmids were designed to perform induced and constitutive co-expression of CCR7 and heterotrimeric G_i_ protein, and the coding genes were synthesized by GenScript for cloning in expression plasmids. The full-length human CCR7 lacking the endogenous signal peptide (UniProt ID: P32248, residues 25-378) was cloned into pcDNA5/TO vector (Thermo Fisher Scientific) with the N-terminal HA secretion signal peptide followed by a FLAG tag, and a C-terminal LgBiT split NanoLuc fragment (32). Single chain human Gγ_2_-Gα_i1_, where Gα_i1_ (UniProt ID: P63096, residues 2-354) with dominant negative mutations (S47N, G203A, E245A, A326S) (31) was fused downstream of Gγ_2_ (UniProt ID: P59768) separated by a 3× GSA linker (30) and cloned into the pEG vector (52). Human Gβ_1_ (UniProt ID: P62873) with C-terminal HiBit NanoLuc complementation fragment and N-terminal 3C protease-cleavable 10×His tag was also cloned into the pEG vector for co-expression.

Protein expression was performed using Expi293Fi cells as per the manufacturer’s recommendation with modifications to use polyethyleneimine (PEI) Max (Polysciences) as a transfection reagent (53), 3 mM sodium valproate and 0.4% (w/v) glucose as enhancers (54), and doxycycline (Wako) as the inducer. The three plasmids were used at an approximately equal weight ratio, and a total of 1 µg plasmid and 4 µg PEI per 1 mL cell culture were mixed in HE400AZ media and incubated for 15 min at room temperature before transfecting the Expi293Fi cells. One day after transfection, the enhancers were added, and after an additional 24 hours at 37°C for initial G_i_ expression, the CCR7 expression was induced by adding 8 µM doxycycline. The culture flask was transferred to a 30°C incubator and shaken for 48 hours. The cells were harvested, washed once with phosphate-buffered saline (PBS), and stored at -30°C. 200 mL Expi293Fi culture typically yields approximately 5 grams of cell pellet.

### Expression and purification of CCL19, CCL21, and scFv16

CCL19 and CCL21 were expressed from Expi293Fi cells but with constitutive expression using a standard CMV promoter, and the coding genes were synthesized by GenScript. Full-length CCL19 (UniProt ID: Q99731) and CCL21 (UniProt ID: O00585) were each fused to the C-terminal 6×His-tagged human IgG1 Fc via the GSGSGSAAA linker, HRV 3C protease site (LEVLFQGP), and GSDGSGSGS linker analogous to the construct previously used to prepare the chemokine domain of CX_3_CL1 (55) but cloned into pD649 transient expression vector (ATUM) (56) instead of the BacMam transfer vector (57). Transfection was performed as described for the CCR7-G_i_ complex with a 1:4 plasmid:PEI ratio, and the enhancers were added to the culture on the following day. The cells were shaken in the 37°C incubator for an additional 84 hours, and the culture supernatant was collected for purification using Ni-NTA (55, 56). The Fc-tagged chemokines were eluted with HEPES-buffered saline (HBS, 10 mM HEPES-Na pH 7.2 and 150 mM NaCl) containing 250 mM imidazole, and the Fc tag was removed by adding 2% (w/w) 3C protease, which was prepared in-house. The solutions were buffer exchanged to HBS containing 10 mM imidazole, flowed through Ni-NTA equilibrated with HBS containing 20 mM imidazole to remove the Fc tag, and concentrated to approximately 300 µL. Due to the limited protein yields (∼50 µM and ∼20 µM for CCL19 and CCL21, respectively), the samples were supplemented with 10% (v/v) glycerol, aliquoted, flash-frozen, and stored at -80°C for further use.

scFv16 (34) was expressed from Hi5 ovarian cells using Bac-to-Bac baculovirus expression system (Thermo Fisher Scientific) and purified as reported previously (55). Briefly, the collected Hi5 culture supernatant was clarified after a precipitate was formed by adding 1 mM NiCl_2_, 5 mM CaCl_2_, and 50 mM Tris-HCl (pH 8), and then incubated with Ni-NTA to capture the C-terminal 6×His-tagged scFv16. The Ni-NTA resin was collected, washed with HBS containing 20 mM imidazole, and eluted by increasing imidazole concentration to 250 mM. The eluate was concentrated and purified using a size-exclusion chromatography (SEC) column Superdex 75 10/300 GL (Cytiva) equilibrated with HBS. The peak fraction was concentrated to approximately 640 µM, aliquoted and flash-frozen with 10% (v/v) glycerol, and stored at -80°C until future use.

### Purification of CCR7-G_i_-scFv16 complexes bound to CCL19 or CCL21

The ligand-free, NanoBiT-tethered CCR7-G_i_-scFv16 complex was purified before the chemokines were added during the final protein concentration step. Approximately 5 grams of cell pellet expressing CCR7-G_i_ was directly solubilized for 2 hours at 4°C with 10 nmol of recombinant scFv16 and nutation in a solubilization buffer consisting of HBS, 10% (v/v) glycerol, 1% (w/v) n-dodecyl-β-D-maltoside (DDM, Anatrace), 0.1% (w/v) lauryl maltose neopentyl glycol (LMNG, Anatrace), 0.11% (w/v) cholesteryl hemisuccinate tris salt (CHS, Anatrace), cOmplete protease inhibitor cocktail (Roche), and apyrase (0.2 µL, New England Biolabs). The resulting cell lysate was clarified by two sequential centrifugations at 12,000 g for 30 minutes at 4°C. The supernatant was then supplemented with 3 mM CaCl_2_ and incubated with 1 mL of in-house anti-FLAG M1 Sepharose for 2 hours at 4°C. The resin was collected in a column and washed sequentially, first with the solubilization buffer, then HBS containing 0.1% LMNG, 0.01% CHS, and 3 mM CaCl_2_. A final wash was performed with HBS containing 0.01% LMNG, 0.001% CHS, and 3 mM CaCl_2_, and the CCR7-G_i_-scFv16 complex was eluted from the column with HBS buffer containing 0.001% LMNG, 0.001% glyco-diosgenin (GDN, Anatrace), 0.0001% CHS, 5 mM EDTA, and 100 µg/mL FLAG peptide (PH Japan). The eluate was concentrated to approximately 400 µL, and 2% (w/w) of in-house 3C protease was added for overnight incubation at 4°C. The protein was subjected to SEC on a Superdex 200 Increase 10/300 GL column (Cytiva) equilibrated with SEC buffer consisting of HBS, 0.001% LMNG, 0.001% GDN, and 0.0001% CHS. Peak fractions were collected, concentrated, and quantified, yielding approximately 0.20 mg (1.1 nmol) of the purified CCR7-G_i_-scFv16 complex.

Concurrently, frozen stocks of the chemokines CCL19 and CCL21 were thawed and dialyzed against the SEC buffer using Xpress Micro Dialyzer with a 3.5 kDa molecular weight cutoff (MWCO) (Scienova GmbH). To form the chemokine-bound signaling complexes, each prepared chemokine was added to the purified CCR7-G_i_-scFv16 in separate reactions. Specifically, CCL19 was added at a 1.6-fold molar excess to 0.6 nmol of CCR7-G_i_-scFv16, while CCL21 was added at the same molar ratio to 0.5 nmol aliquot of the ligand-free complex. Finally, each mixture was concentrated to 4.5 mg/mL using 3 kDa MWCO Amicon Ultra 0.5 mL centrifugal filters for cryo-EM.

### Cryo-EM grid preparation and data collection

For cryo-EM analysis, a 3.5 µL aliquot of either complex was applied to a glow-discharged Quantifoil R1.2/1.3 300-mesh gold grid. The grids were plunge-frozen in liquid ethane using a Vitrobot Mark IV (Thermo Fisher Scientific) with a 10-sec hold period, blot force of 10, and blotting time of 2-4 sec with 100% humidity at 4°C.

Cryo-EM data were subsequently collected using a JEOL JEM-Z320FHC electron microscope operated at 300 kV, equipped with an in-column omega energy filter set to a 20 eV slit width. Images were recorded on a Gatan K3 camera in correlated double sampling mode, with a calibrated pixel size of 0.79 Å at the specimen. Automated data acquisition was performed using SerialEM (58) with a 3 × 3 image-shift pattern and a nominal defocus range of -1.0 to -1.8 μm. Each movie was recorded for 2.6 seconds, fractionated over 50 frames, yielding a total exposure of 50 electrons/Å^2^ and an exposure rate of 1 electron/Å^2^ per frame. A total of 20,142 movies (7,506 + 12,636) were collected from two grids for the CCL19-CCR7-G_i_-scFv16 complex, while 5,968 movies (2,376 + 3,592) were collected from two grids for the CCL21-CCR7-G_i_-scFv16 complex.

### Cryo-EM data processing

The data were analyzed using cryoSPARC version 4 (59) for both signaling complexes unless otherwise noted, and the data processing schemes are summarized in SI Appendix, Fig. S3 and S4. Patch motion correction and contrast transfer function (CTF) estimation were performed. Particles were picked iteratively with template-based picking using a representative GPCR-G protein complex and Topaz picking (60) combined with 2D classifications.

For the CCL19-CCR7-G_i_-scFv16 complex (SI Appendix, Fig. S3), 3,374,958 initial particles were extracted and subjected to Non-uniform Refinement (NU-refine) with Global and Local CTF Refinements. A representative GPCR-G protein-scFv16 complex was used as an initial reference map low-pass filtered to 30 Å. 3D classification was first performed to eliminate the complex particles with poor 7TM density. Several rounds of Heterogeneous Refinement (Hetero-Refine) and NU-Refine were performed to isolate a homogeneous subset of particles. We then utilized 3D Variability Analysis (3DVA) followed by Hetero-Refine using the resulting 3DVA volumes as references. While we observed heterogeneity related to glycosylation, such as at N292, we prioritized classes with the best-defined chemokine core features. The resulting class was further refined using Reference-Based Motion Correction (RBMC), NU-refine, and Local CTF Refinement. The selected 256,195 particles were subjected to the final NU-Refine, which yielded a map with a nominal resolution of 3.0 Å as determined by the gold-standard Fourier shell correlation (GSFSC) at the 0.143 threshold.

For the CCL21-CCR7-G_i_-scFv16 complex (SI Appendix, Fig. S4), 1,537,777 initial particles were processed similarly. After initial 2D and 3D classification, particles representing the best class and minor views from the second-best class were merged and subjected to iterative rounds of NU-Refine, RBMC, and Local CTF-Refine. Subsequent 3DVA-based Hetero-Refine steps were employed to address conformational heterogeneity at the chemokine core. Notably, we identified minor classes exhibiting weak or partial density corresponding to potential N-linked glycosylation at N36 and N292 of CCR7, as well as ambiguous density near the extracellular surface potentially representing parts of the flexible CCL21 C-terminal tail (SI Appendix, Fig. S4A). However, these classes constituted a small fraction of the dataset and displayed lower homogeneity and resolution in the core chemokine-receptor interface compared to the major population. Therefore, we focused on the most homogeneous subset to achieve the highest local resolution (quality or modelability) of the chemokine region. After several rounds of refinement and classification, the final set of 195,011 particles yielded a map with a nominal resolution of 3.2 Å determined by GSFSC at 0.143 threshold.

The maps were sharpened with a B-factor of -60 Å^2^ for model building, refinement and visualization of the map-model correlation. For both datasets, local resolution was estimated using cryoSPARC, and the maps were further sharpened using deepEMhancer for overall map presentation purposes only (61). UCSF pyEM v0.5 (62) was used to plot particle viewing directions on the 3D maps.

### Model building and refinement

The initial models were prepared using AlphaFold 3 (63), and each subunit was roughly fitted into the map by real-space rigid body refinement using Phenix (64). The models were subjected to iterative rounds of manual rebuilding in Coot (65) and real-space refinement using Phenix. Secondary structure restraints and geometry restraints were applied during refinement. The map-model correlation and refinement statistics are summarized in SI Appendix, Table S1 and Fig. S6. Figures were prepared using UCSF ChimeraX (66) and PyMOL (Schrödinger, LLC).

### Molecular dynamics simulations

Input models and parameters for all-atom MD simulations of CCR7 complexed with CCL19 or CCL21 were generated using CHARMM-GUI (67–69). All molecules other than CCR7 and CCL19/21 were stripped from the cryo-EM models. The unmodelled sidechains were manually built and refined with Coot (65). The internal water molecules were placed based on the estimation by HOMOLWAT (70). The unmodelled part of the N-terminus and the C-terminus of CCR7 and CCL19/21 were left unmodelled, and the terminal amino groups and the carboxyl groups were neutrally capped by acetylation (ACE patch) and methylamidation (CT3 patch), respectively. Protonation states of the titratable residues at pH 7.0 were determined with PROPKA (71). The terminal amino groups of the native N-terminus of CCL19 (G1) and CCL21 (S1) were deprotonated by the NGNE and NNEU patches, respectively (estimated pKa 5.96 and 4.51). Because the NGNE patch was not available in CHARMM-GUI, a local install of CHARMM c49b2 (72) was used for the patching using the scripts generated by CHARMM-GUI. All the aspartic acid and glutamic acid residues were negatively charged except D104^2.50^ (estimated pKa 7.50 in CCR7-CCL19 and 7.81 in CCR7-CCL21). All the lysine and arginine residues were positively charged. All the histidine residues were neutralized by δ-nitrogen protonation. The model was embedded in a rectangular lipid bilayer comprising 203 POPC molecules (102 in the upper leaflet and 101 in the lower leaflet) with the orientation estimated by PPM2.0 (73). The system was solvated and charge-neutralized by ∼24,000 water molecules (water layers of 22.5 Å thickness on top and bottom of the proteins) and 150⍰mM NaCl. The system dimensions and the total number of atoms were 90⍰Å⍰× ⍰90⍰Å⍰× ⍰141⍰Å and 107,178 atoms for CCL19-CCR7, 90⍰Å⍰×190⍰Å⍰× ⍰139⍰Å and 105,680 atoms for CCL21-CCR7 (SI Appendix, Fig. S9A). The CHARMM36m force-field parameters (74) were used for the proteins, lipids, and ions. The TIP3P model was used for water.

MD simulations were performed using GROMACS 2024.3 (75). Each system was energy-minimized using the steepest-descent algorithm for 10,000 steps. Then, six independent MD runs (replicas) were performed. For each replica, the system was equilibrated and relaxed in the following way. First, temperature was elevated from 0 to 100 K for 20 ps in the NVT ensemble using the V-rescale thermostat (76). Temperature was further elevated from 100 K to 310.15 K under the pressure of 1 bar for 400 ps in the NPT ensemble using the V-rescale thermostat and the Berendsen barostat (77). During the energy minimization and the temperature elevation, the x, y, z positions of the backbone heavy atoms, the x, y, z positions of the sidechain heavy atoms, the z positions of the POPC phosphorous atoms, the dihedral angles about the double bond in the POPC alkyl chain, were harmonically restrained to the initial configuration at the force constant of 4000 kJ/mol/nm^2^, 2000 kJ/mol/nm^2^, 1000 kJ/mol/nm^2^ and 1000 kJ/mol/rad^2^, respectively. Then the restraints were gradually relaxed towards 0 for 35 ns in the NPT ensemble of 310.15 K and 1 bar using the V-rescale thermostat and the C-rescale barostat (78). After that, the production run was performed for 2.3 μs in the NPT ensemble of 310.15 K and 1 bar using the V-rescale thermostat and the C-rescale barostat at the timestep of 2 fs. Bond lengths involving hydrogen atoms were constrained using the LINCS algorithm (79). Long-range electrostatic interactions were calculated with the particle mesh Ewald method (80). Atomic coordinates were saved every 100 ps. The simulations were carried out using the TSUBAME4.0 supercomputer at Institute of Science Tokyo.

The molecules were made ‘whole’ and put back into the simulation cell using the gmx trjconv program in the GROMACS package. The trajectory analysis was performed using MDAnalysis v2.9.0 (81) and the graphs were visualized using Matplotlib (82). To align the trajectories, the corresponding cryo-EM models were used as the alignment references, which were transformed so that the (x, y) origin is the center of the proteins and the z origin is the center of the membrane, and the z-axis is perpendicular to the membrane (SI Appendix, Fig. S10B).

In all the trajectory analysis except the ICL1 conformational analysis (SI Appendix, Fig. S12D), the trajectories were least-squares aligned to the reference models by the Cα atoms of CCR7 except the flexible N-terminal (−50) and ECL2 (200-209) regions. In the ICL1 conformational analysis (SI Appendix, Fig. S12D), the trajectories were aligned by the Cα atoms of the ICL1-flanking residues of TM1 and TM2 (78-85, 92-99).

In all the trajectory analysis except the simulation time course plots, the first 1 μs of the 2.3 μs simulation were excluded from the analysis as the relaxation (equilibration) period, because the average RMSD (SI Appendix, Fig. S9B) and the TM displacements (SI Appendix, Fig. S10A) were rapidly changing in this period.

The MD snapshots used in the figures were selected so that the TM centroid coordinates and the H8 orientation (the angle and the TM2-G330 distance) are closest to the MD average (the timestep 1597.8 ns of replica #1 for CCL19-CCR7, the timestep 1236.1 ns of replica #1 for CCL21-CCR7).

The density plots in Fig. 5A,B and in SI Appendix, Fig. S12A, B were calculated using the Gaussian kernel density estimation implemented in Scikit-learn (83).

### Plasmids for signaling and expression analysis

The cDNAs encoding full-length human CCR7 (WT) and four point mutants (Y83A, R88A, D94A, and D336A) were chemically synthesized and cloned into the pcDNA3.1(+) vector by GenScript. To facilitate cell surface expression and detection, an HA signal peptide sequence followed by a FLAG epitope tag was genetically fused to the N-terminus of each receptor construct.

### Cellular CCR7 signaling

#### Cell culture and transfection

HEK293T cells (RIKEN BRC) were seeded at 0.8 × 10^6^ cells per well in a 6-well plate for G protein dissociation assay, and 1.6–1.8 × 10^6^ cells per 6-cm dish for β-arrestin recruitment assay in Dulbecco’s modified Eagle’s medium (DMEM, 1 g/L glucose; Wako) supplemented with 10% (v/v) fetal bovine serum (FBS; Sigma-Aldrich), 100 U/mL penicillin, and 100 µg/mL streptomycin (Nacalai Tesque) one day before transfection. Cells were incubated at 37°C in a humidified atmosphere with 5% CO_2_.

For the NanoBiT G protein dissociation assay, cells were transfected with 200 ng WT or mutant CCR7 in pcDNA3.1, together with 100 ng Gα_i1_–LgBiT in pcDNA3.1, 500 ng Gβ_1_ in pcDNA3.1, 500 ng SmBiT–Gγ_2_(C68S) in pCAGGS, and 100 ng Ric8A in pCAGGS. Plasmids were diluted in Opti-MEM I (Thermo Fisher Scientific) and mixed with 5 µg PEI MAX, incubated for 15 min at room temperature, and added to the cells.

For the NanoBiT β-arrestin recruitment assay, cells were transfected with 400 ng WT or mutant CCR7 in pcDNA3.1, together with 200 ng SmBiT–β-arrestin2 in pCAGGS, and 1 µg LgBiT–CAAX in pCAGGS. Plasmids were diluted in Opti-MEM I and mixed with 10 µg PEI MAX, incubated for 15 min at room temperature, and added to the cells.

#### NanoBiT G protein dissociation assay

24 hours after transfection, cells were washed with PBS, detached with 0.53 mM EDTA in PBS, diluted with Hank’s balanced salt solution (HBSS, pH 7.2; Gibco), and centrifuged at 100 × *g* for 3 min. After removal of the supernatant, cells were resuspended in 2 mL of HBSS containing 0.01% (w/v) fatty acid-free bovine serum albumin (BSA; Nacalai Tesque) as assay buffer. The cell suspension was dispensed into a white 384-well plate at 24 µL per well, followed by addition of 6 µL of 50 µM coelenterazine (Selleck Chemicals), diluted in assay buffer. After incubation for 2 h at room temperature, baseline luminescence was recorded using a SpectraMax Mini plate reader (Molecular Devices). Thereafter, 6 µL of 6× ligands prepared in assay buffer were manually added to each well. CCL19 (BioLegend, Cat# 582106) and CCL21 (BioLegend, Cat# 582208) were initially prepared at 1 µM and 1.39 µM, respectively, and serially diluted in assay buffer in fivefold steps. The plate was immediately read in kinetic mode, and luminescence counts were recorded for 25 min with an integration time of 0.5 sec per read at 2-min intervals. The luminescence signal remained stable between 8 and 25 min, and data obtained between 10 and 22 min (six time points) were used for analysis.

#### NanoBiT β-arrestin recruitment assay

24 hours after transfection, cells were harvested as described above, and resuspended in 4 mL of assay buffer. The cell suspension was dispensed into a white 96-well plate at 80 µL per well, followed by addition of 20 µL of 50 µM coelenterazine diluted in assay buffer, and baseline luminescence was recorded. Thereafter, 20 µL of 6× ligands prepared as described above were manually added to each well, and luminescence counts were recorded for 20 min with an integration time of 0.5 sec per read at 1-min intervals. The luminescence signal remained stable between 4 and 20 min, and data obtained between 5 and 12 min (six time points) were used for analysis.

#### Data analysis

The baseline luminescence recorded prior to ligand addition was used for normalization. For each well, the mean of six consecutive luminescence values within the indicated time window was calculated, and the resulting value was expressed as a fold change relative to the baseline for subsequent analysis. Concentration–response curves were analyzed using Prism 10 (GraphPad Software) by nonlinear regression with a variable slope (four-parameter logistic) model. Independent experiments (n = 3) were performed, with WT and mutant receptors analyzed in parallel within each experiment. For each experiment, EC_50_ values and E_max_ of G_i1_ dissociation or β-arrestin2 recruitment were calculated. Statistical comparisons were performed using a paired, two-tailed Student’s t-test for WT versus Y83A. For comparisons among WT, R88A, D94A, and D336A, a one-way repeated-measures ANOVA followed by Dunnett’s multiple comparisons test (WT vs. each mutant) was applied. A *P* value < 0.05 was considered statistically significant.

### Cell surface CCR7 expression

#### Cell culture and transfection

HEK293F cells were maintained in suspension in FreeStyle 293 Expression Medium (Thermo Fisher Scientific) at densities ranging between 0.5 × 10^6^/mL and 2 × 10^6^/mL and shaken at 37°C and 130 rpm with 8% (v/v) CO_2_. For transfection, cells were prepared at a density of 1.0 × 10^6^/mL in a 1 mL volume. Transfection complexes were prepared by mixing 1 µg of plasmid DNA and 4 µg of PEI STAR transfection reagent (Tocris Bioscience) in 100 µL of Opti-MEM (Thermo Fisher Scientific). The mixture was incubated for 10 min at room temperature and added to the cell suspension. The empty pcDNA3.1(+) vector was used as a negative control. Cells were cultured for 24 h prior to analysis.

#### Flow cytometry

24 hours post-transfection, cells were harvested and washed twice with 1 mL of ice-cold PBE buffer [DPBS (Nacalai Tesque) supplemented with 0.1% BSA and 1 mM EDTA] by centrifugation at 300 × *g* for 3 min at 4°C. The cell pellets were resuspended in 100 µL of PBE buffer containing FITC-conjugated anti-FLAG M2 monoclonal antibody (Sigma-Aldrich, Cat# F4049, 1:100 dilution) and incubated for 1 hour at 4°C. Following incubation, cells were washed twice with 1 mL of PBE and resuspended in 500 µL of the same buffer. The cell surface expression levels were analyzed using a FACSLyric flow cytometer (BD Biosciences). Data collection and analysis were performed using BD FACSuite software. FITC fluorescence was excited by the 488 nm blue laser, separated by a 507 nm long-pass mirror, and detected through a 527/32 nm bandpass filter. A sequential gating strategy was applied to isolate single cells: first, cell debris was excluded based on forward scatter (FSC-A) and side scatter (SSC-A) characteristics. Subsequently, singlets were selected by excluding doublets using FSC-H versus FSC-W and SSC-H versus SSC-W plots. The mean fluorescence intensity (MFI) of the FITC signal in the singlet population was quantified. Statistical analysis was performed on the unnormalized raw MFI data using a one-way ANOVA followed by Dunnett’s multiple comparisons test against the WT group in Prism 8 (GraphPad Software).

## Supporting information

This PDF file includes: Figures S1 to S12 and Table S1

## Data, Materials, and Software Availability

The cryo-EM maps and the corresponding atomic coordinates have been deposited in the Electron Microscopy Data Bank (EMDB) and the Protein Data Bank (PDB) under accession codes EMD-66874 (PDB: 9XHH) and EMD-66875 (PDB: 9XHI) for CCL19-CCR7-G_i_-scFv16 and CCL21-CCR7-G_i_-scFv16, respectively. The MD simulation trajectories generated in this study are available at Zenodo [https://doi.org/10.5281/zenodo.18859194]. Any additional information required to reanalyze the data reported in this paper, as well as unique and stable reagents generated in this study, are available from the corresponding author upon reasonable request.

## Acknowledgements

We thank Gaku Nakamura and Dr. Asuka Inoue for providing the NanoBiT assay plasmids and technical guidance. We also thank the members of the Cellular and Structural Physiology Laboratory (CeSPL) for their daily scientific discussions, and Ms. Miho Sasaki for her research support. We used Google Gemini (Gemini 3.1 Pro) to edit the text and enhance readability, maintaining full oversight, fact-checking all outputs, and taking full accountability for the final content. This work was supported by the Japan Society for the Promotion of Science (JSPS) KAKENHI (grant numbers JP24K01965 and JP24K21935 to N.T., and JP20H00451 to Y.F.), the Kobayashi Foundation (grant number 303 to Y.S.), and the Kurume University Ishibashi Foundation for the Promotion of Science (to Y.S.). The MD simulations were performed using the TSUBAME4.0 supercomputer at the Institute of Science Tokyo.

