## Supplementary material for "Structural Insights into Biased Signaling at Chemokine Receptor CCR7": This PDF file includes: Figures S1 to S12 and Table S1

Naotaka Tsutsumi*

**This PDF file includes:**

**Figures S1 to S12**

**Table S1**

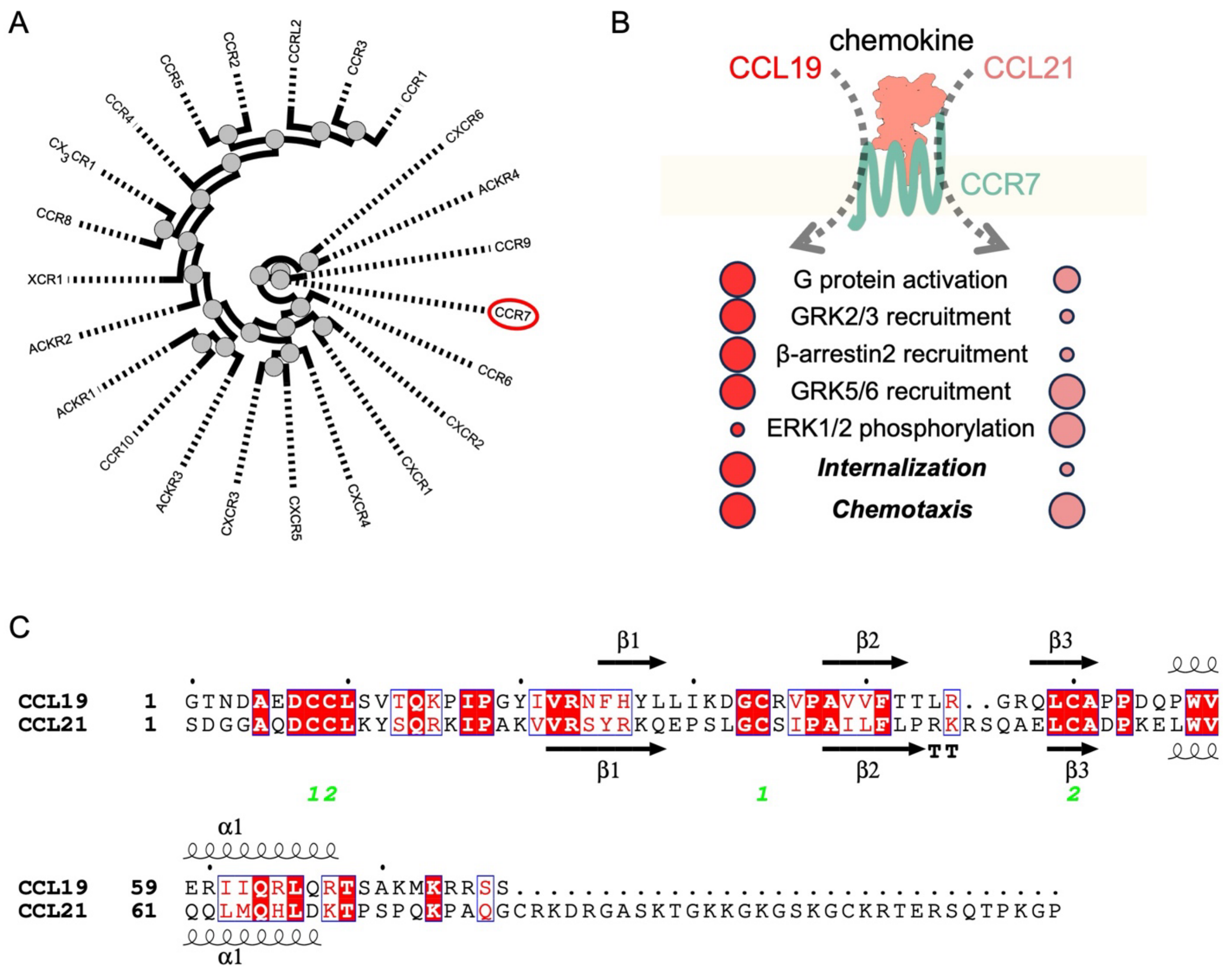


**Fig. S1 | Chemokine receptors and CCR7-binding chemokines**

(A) Phylogenetic tree of human chemokine receptors rooted on CCR7. The tree was generated with the sequences at structurally conserved (generic) positions of the receptors using the GPCRdb website (1). (B) Comparison of signaling and functional profiles of the CCR7 axis activated by CCL19 (left) and CCL21 (right) based on the literature (2–7). The dashed arrows indicate the activation flow. Circle size schematically represents the relative magnitude of signaling activation or functional outcomes. (C) Sequence alignment of human CCL19 and CCL21. Secondary structure elements are indicated on the top and bottom of the sequences. CCL21 possesses a long, basic C-terminal tail, which was not clearly visualized in the current structure.

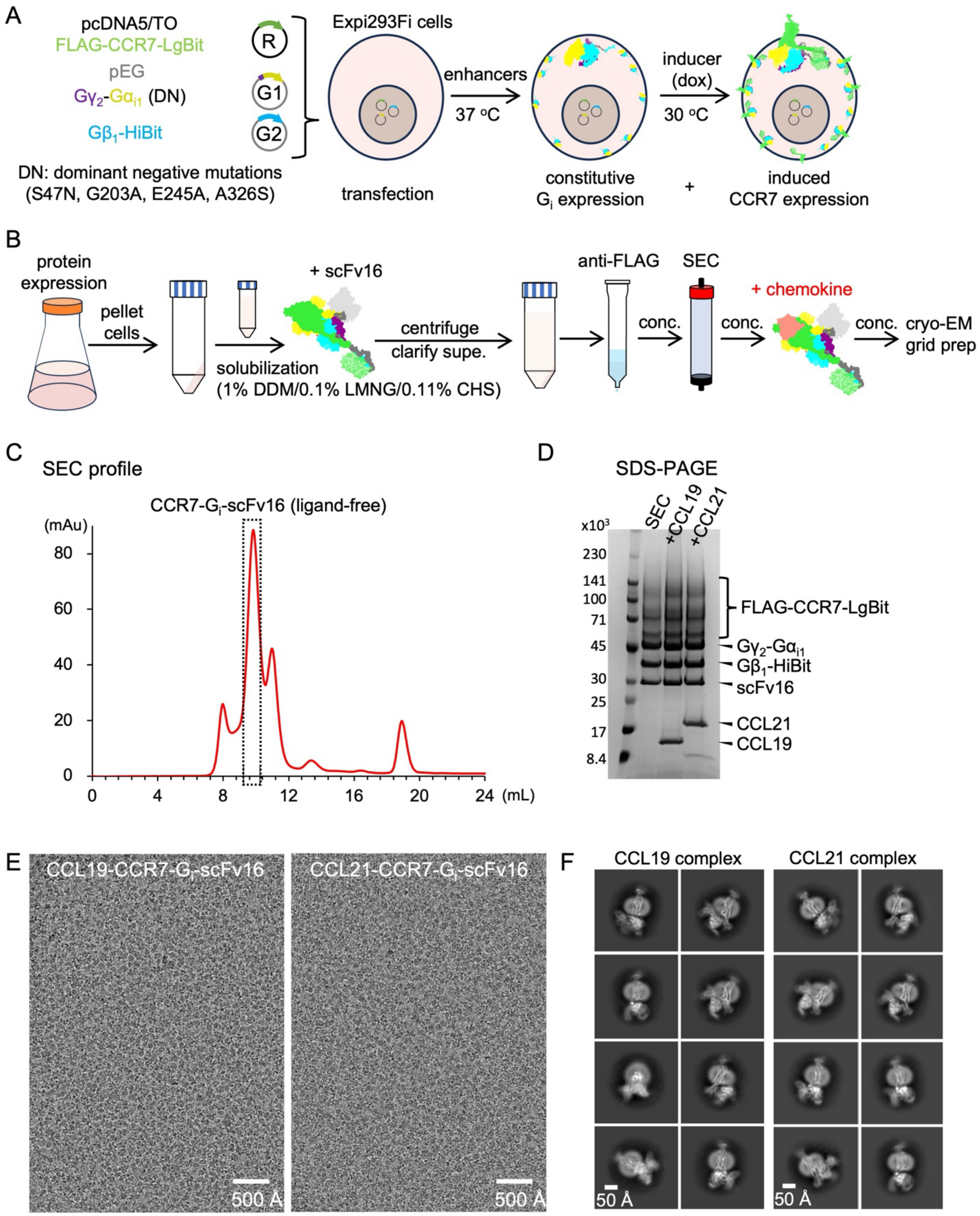


*Fig. S2 caption on the next page.*

**Fig. S2 | Temporally controlled mammalian expression of the CCR7-$G_i$ complexes for cryo-EM analysis.**

(A) Schematic representation of the temporally controlled expression system in inducible Expi293F cells. Plasmids encoding FLAG-CCR7-LgBiT (R; inducible, pcDNA5/TO) and $G_i$ heterotrimer components [constitutive, pEG; Gγ2-G$\alpha_{i1}$(DN) (G1) and G$\beta_1$-HiBiT (G2)] are co-transfected. Cells are cultured at 37°C for constitutive $G_i$ expression, followed by induction with doxycycline (dox) and temperature shift to 30°C for induced CCR7 expression and sustained $G_i$ expression. DN denotes dominant negative mutations (S47N, G203A, E245A, A326S) in G$\alpha_{i1}$. (B) Workflow for the purification of the CCR7-$G_i$-scFv16 complexes, highlighting the strategy of purifying the apo-complex followed by the addition of chemokines. (C) Size-exclusion chromatography (SEC) profile of the ligand-free CCR7-$G_i$-scFv16 complex. The peak fraction used for subsequent steps is indicated by the dashed box. Purified chemokines are added to this sample for cryo-EM analysis. (D) SDS-PAGE analysis of the SEC peak fraction and the final complexes after the addition of CCL19 or CCL21. (E) Representative cryo-EM micrographs (scale bar 500 Å) of the CCL19-CCR7-$G_i$-scFv16 complex (left) and the CCL21-CCR7-$G_i$-scFv16 complex (right). (F) Selected 2D class averages (scale bar 50 Å) of the CCL19-CCR7-$G_i$-scFv16 complex (left) and the CCL21-CCR7-$G_i$-scFv16 complex (right).

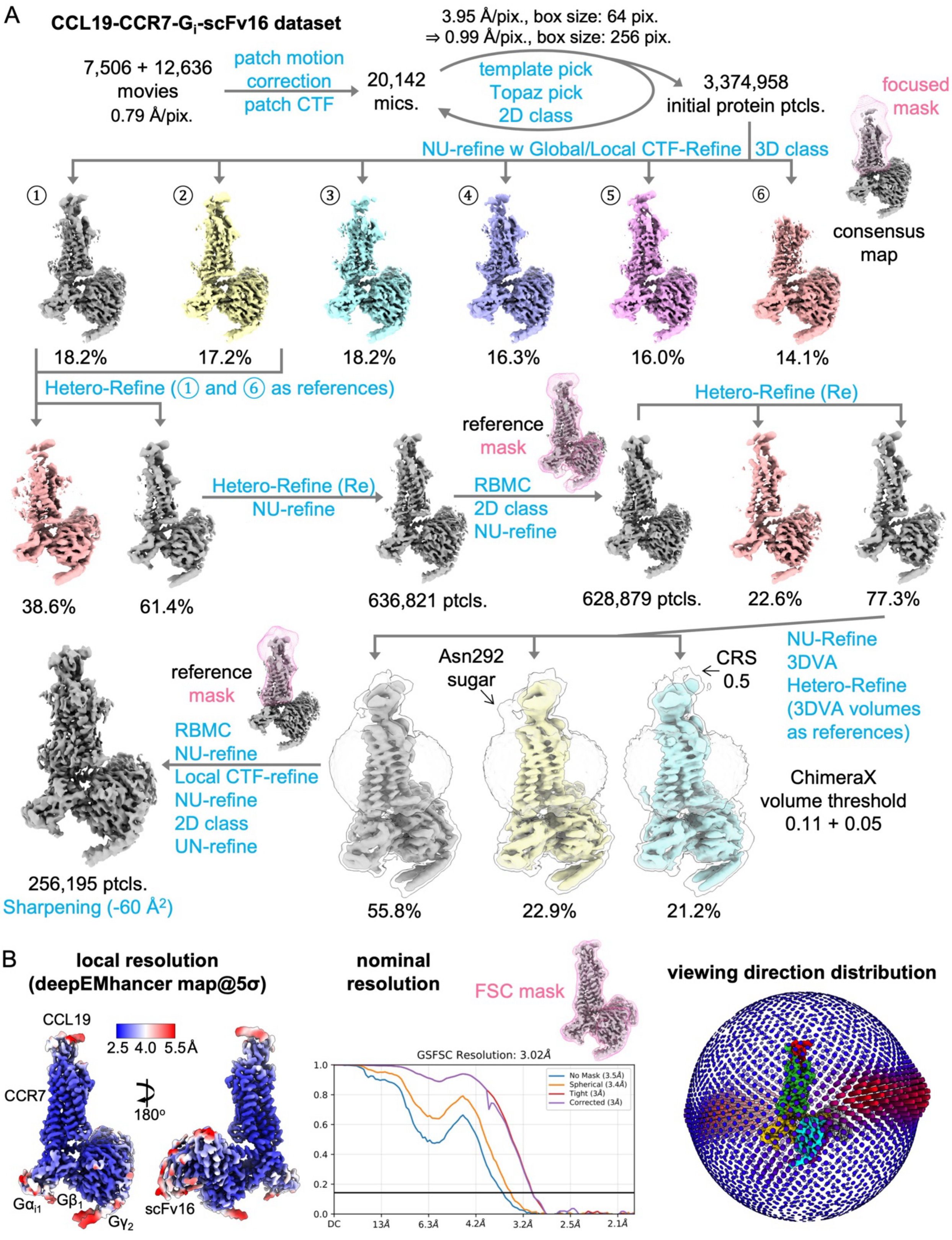


*Fig. S3 caption on the next page.*

**Fig. S3 | Cryo-EM data processing workflow for the CCL19-CCR7-$G_i$-scFv16 complex.**

(A) Schematic overview of the cryo-EM data processing pipeline in cryoSPARC (8). The workflow involved iterative classification and refinement strategies to address conformational heterogeneity, particularly concerning the CCR7 and chemokine regions. Patch CTF: Patch CTF Estimation, template pick: Template Picker, Topaz pick: training and extraction by Topaz, 2D class: 2D Classification, NU-Refine: Non-uniform Refinement, CTF-Refine: CTF Refinement, 3D class: 3D Classification, Hetero-Refine: Heterogeneous Refinement, RBMC: Reference-Based Motion Correction, 3DVA: 3D Variability Analysis. (B) Validation of the final cryo-EM reconstruction. Left: Local resolution estimation colored on the deepEMhancer-sharpened map at 5σ contour level, according to the scale bar. Center: Gold-standard Fourier shell correlation (GSFSC) curves indicating an overall nominal resolution of 3.0 Å (FSC=0.143 criterion). Right: Viewing direction distribution shown by 3D histograms overlaid on the 3D map. The height and color from blue to red indicate the relative number of particles from the specific direction.

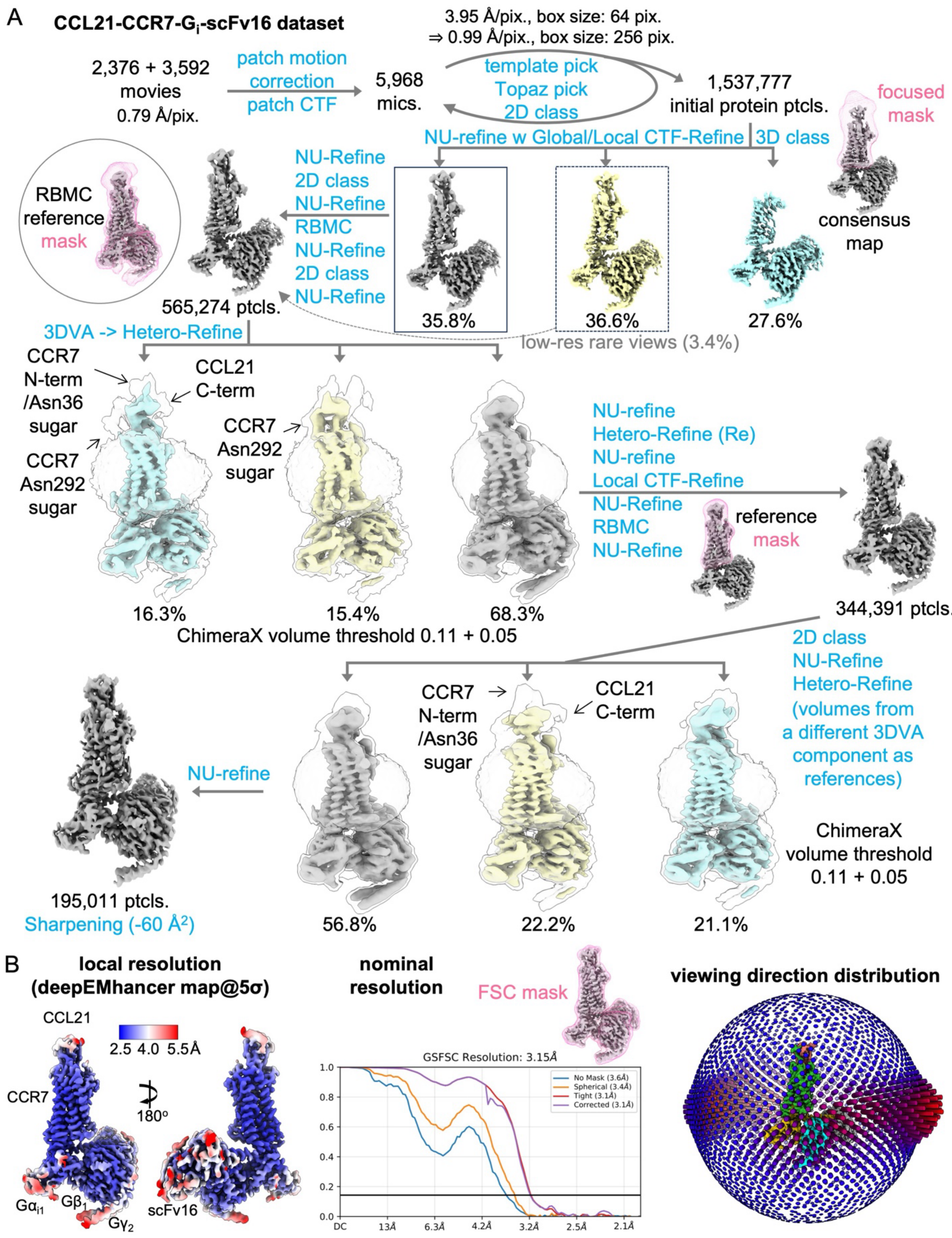


*Fig. S4 caption on the next page.*

**Fig. S4 | Cryo-EM data processing workflow for the CCL21-CCR7-$G_i$-scFv16 complex.**

(A) Schematic overview of the cryo-EM data processing pipeline for the CCL21 complex. The workflow involved iterative classification and refinement strategies to address conformational heterogeneity, particularly concerning the CCR7 and chemokine regions. (B) Validation of the final cryo-EM reconstruction. Left: Local resolution estimation colored on the deepEMhancer-sharpened map at 5σ contour level, according to the scale bar. Center: GSFSC curves indicating an overall nominal resolution of 3.2 Å (FSC=0.143 criterion). Right: Viewing direction distribution shown by 3D histograms overlaid on the 3D map. The height and color from blue to red indicate the relative number of particles from the specific direction.

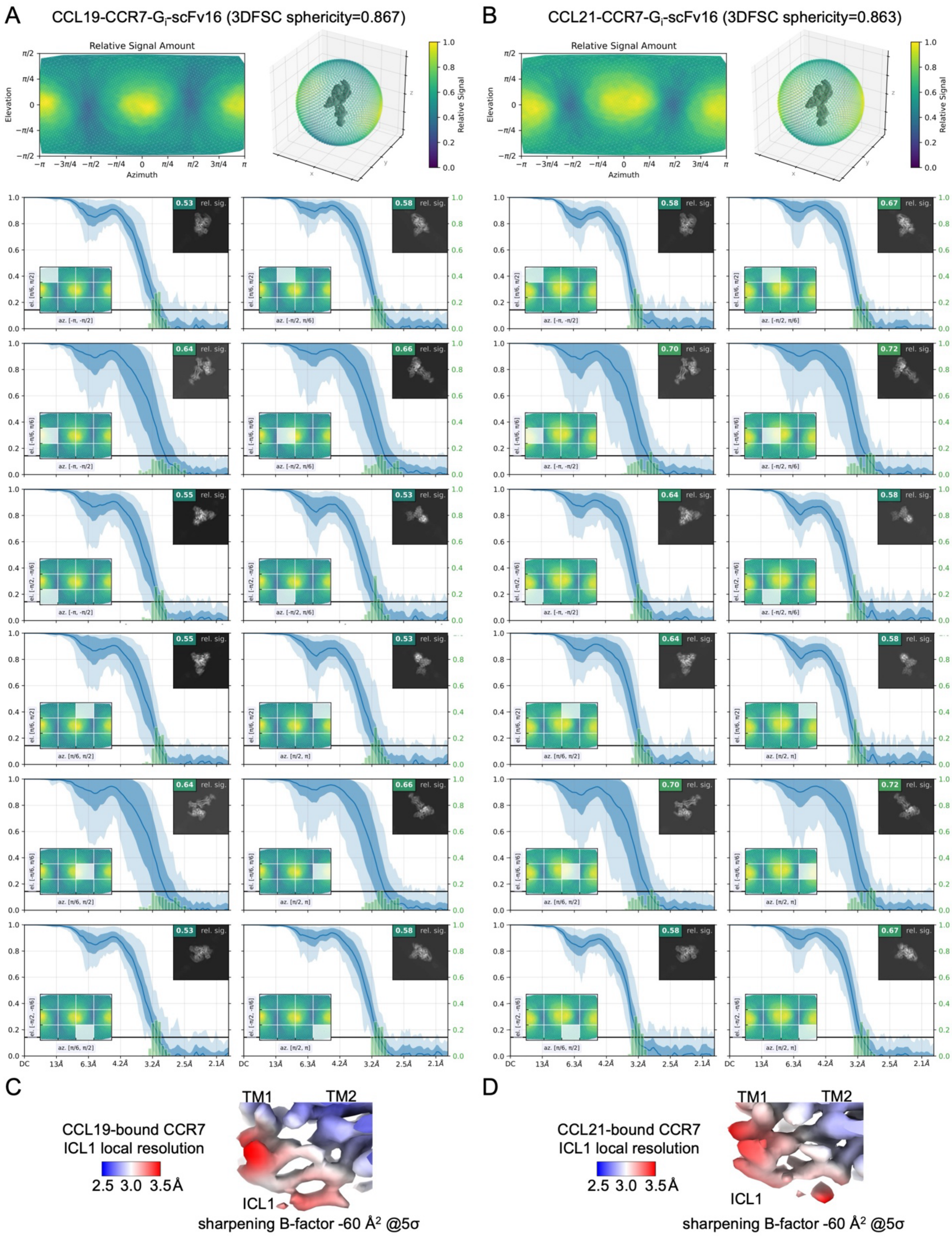


*Fig. S5 caption on the next page.*

**Fig. S5 | Orientation diagnostics and local resolution of the CCR7 complexes.**

Orientation diagnostics for (A) CCL19-CCR7-$G_i$-scFv16 and (B) CCL21-CCR7-$G_i$-scFv16 generated by cryoSPARC. The sphericity values calculated using 3DFSC (9) are 0.867 and 0.863, respectively. In blue: statistics over cFSC curves: mean, min, max, ± one standard deviation plotted against spatial frequency. In green: histogram over 0.143 crossings of the same curves. Local resolution maps of the intracellular loop 1 (ICL1) region for (C) CCL19-bound and (D) CCL21-bound CCR7. Unlike ICL2 and ICL3, ICL1 is not involved in the interaction with $G_i$, resulting in a relatively lower map quality in this region, implying a higher degree of local flexibility. Therefore, the structural model built from the map represents the most probable conformation of ICL1.

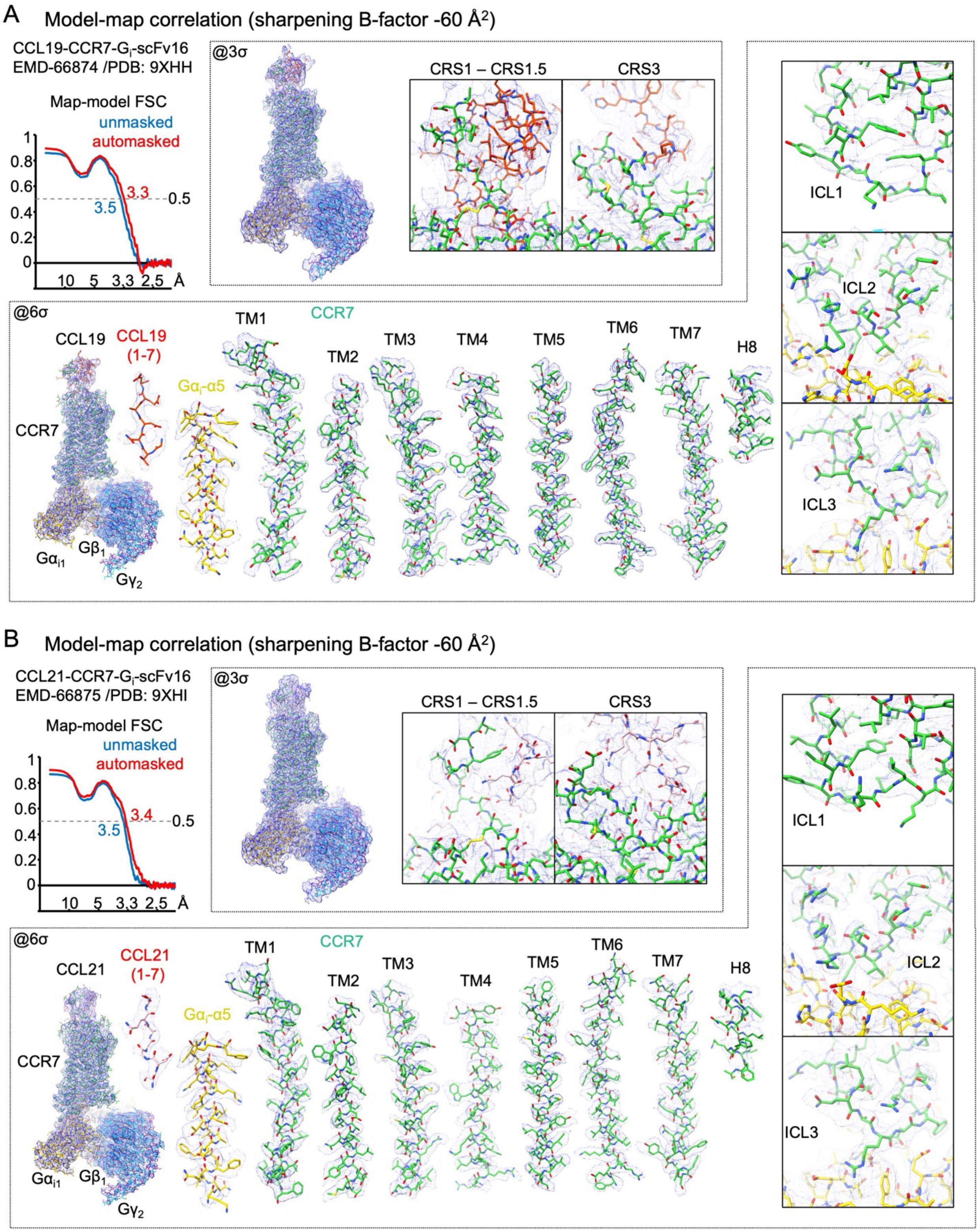


*Fig. S6 caption on the next page.*

**Fig. S6 | Model-map correlation and representative cryo-EM densities.**

(A) Validation of the CCL19-CCR7-$G_i$-scFv16 structural model. Top left: Map-model FSC curves. Top center: Overview and close-up views of the chemokine-receptor interface (CRS1-CRS1.5 and CRS3). The model is overlaid with the cryo-EM density map (blue mesh) contoured at 3σ. Bottom and right: Overview of the map contoured at 6σ (left) and representative densities for CCR7 TM helices (TM1-7), Helix 8 (H8), CCL19 N-terminus (residues 1-7), the $G\alpha_i$ α5 helix, and ICL1-3 contoured at 6σ. (B) Validation of the CCL21-CCR7-$G_i$-scFv16 structural model, presented similarly to panel A.

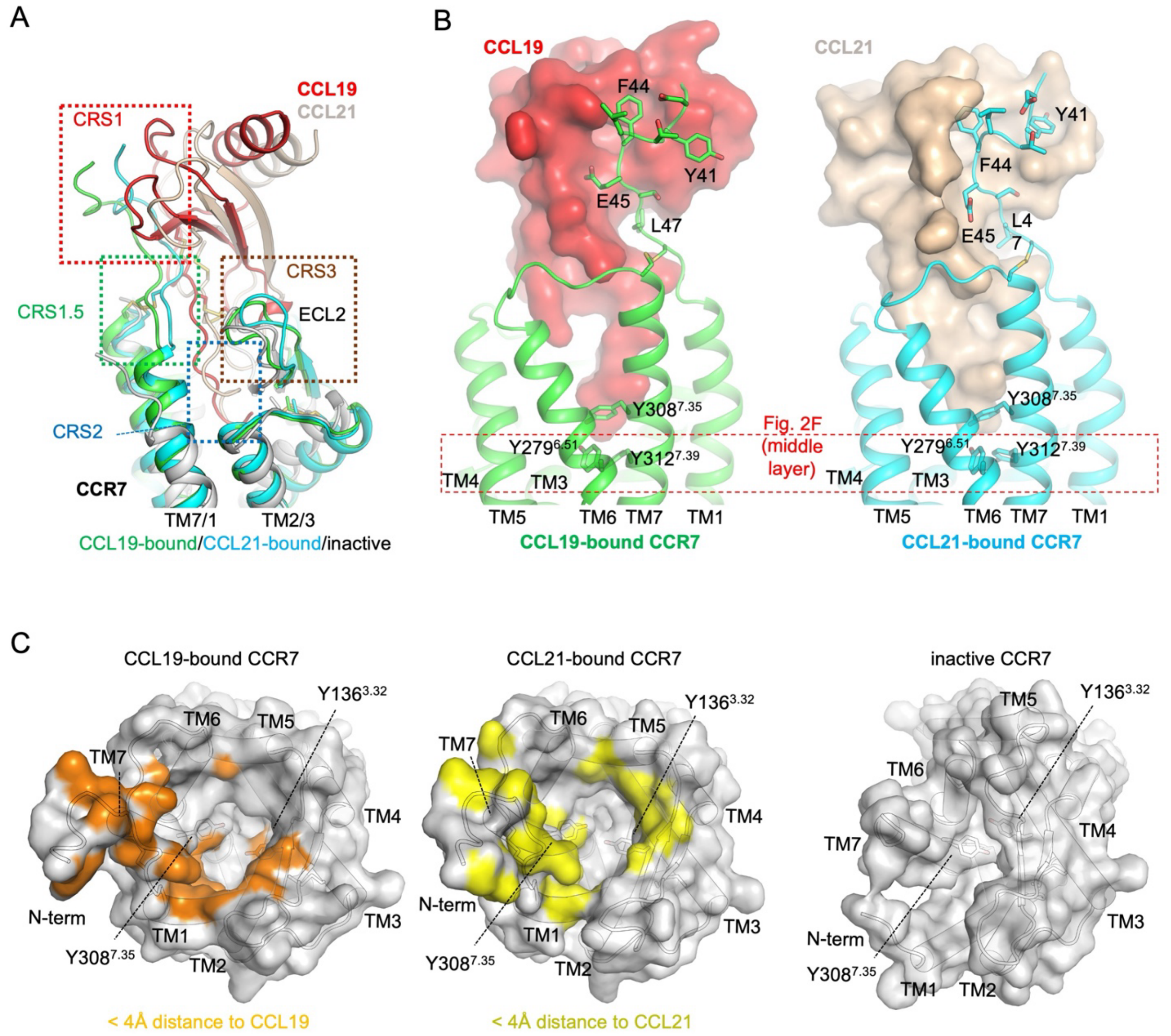


**Fig. S7 | Chemokine recognition interfaces and binding footprints on CCR7.**

(A) Overview of the chemokine recognition sites. Dashed boxes indicate CRS1, CRS1.5, CRS2, and CRS3. (B) Global views of the CRS1 interface and tyrosine residues at CRS2. CCL19 (left, red) and CCL21 (right, orange) are shown in surface representation interacting with CCR7 (green or cyan, depicted as ribbons and sticks). Key CCR7 residues (Y41, F44, E45, and L47) involved in the CRS1/CRS1.5 interactions are highlighted as sticks. The boxed region corresponds to the middle-layer aromatic cluster shown in Fig. 2F. (C) Chemokine footprints within the CCR7 binding pocket. Top views of the CCR7 surface are colored based on their distance to CCL19 (left) or CCL21 (center), with residues within 4 Å of the respective chemokines highlighted. The inactive CCR7 structure (right) is shown for comparison to highlight its closed binding pocket conformation. The positions of key aromatic residues (Y136$^{3.32}$ and Y308$^{7.35}$) are indicated for reference.

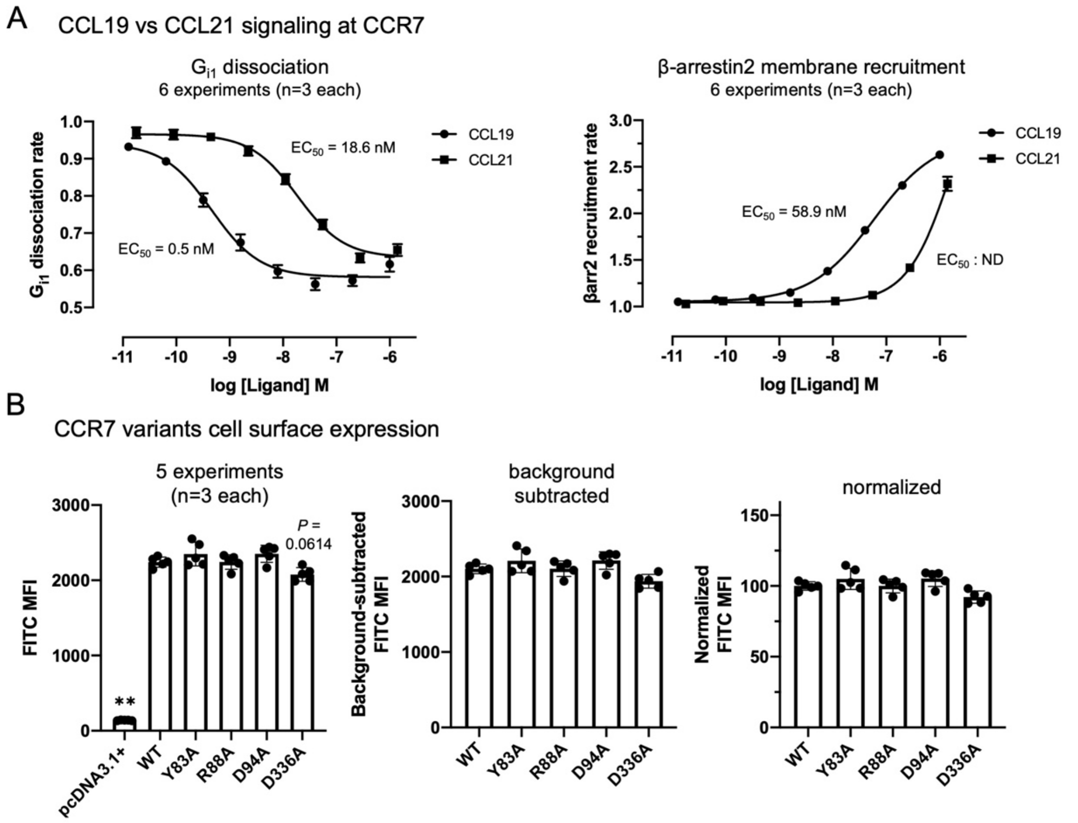


**Fig. S8 | Signaling profiles of CCL19 and CCL21 at CCR7 and cell surface expression of CCR7 variants.**

(A) CCL19 and CCL21 induce $G_{i1}$ dissociation and β-arrestin2 recruitment at WT CCR7. Concentration-response curves for $G_{i1}$ dissociation rate (left) and β-arrestin2 membrane recruitment rate (right). Curves were fitted using a four-parameter logistic model, and the calculated $EC_{50}$ values are indicated on the graphs. In the $G_{i1}$ dissociation assay, CCL19 (~0.5 nM) was ~40-fold more potent than CCL21 (~18.6 nM), though both achieved comparable maximal efficacy. In the β-arrestin2 recruitment assay, CCL19 induced a robust response ($EC_{50}$ ~58.9 nM), whereas the substantially lower potency of CCL21 precluded accurate estimation of its $EC_{50}$ and efficacy (not determined, ND). Data represent mean ± SEM of 6 independent experiments, each performed in triplicate. (B) Cell surface expression of CCR7 variants. Flow cytometry analysis assessing the cell surface expression of WT CCR7 and the indicated mutants (Y83A, R88A, D94A, D336A). Cells transfected with the empty vector (pcDNA3.1+) were used as a negative control to determine background fluorescence. Bar graphs show raw FITC mean fluorescence intensity (MFI) (left), background-subtracted FITC MFI (middle), and normalized FITC MFI relative to WT expression levels (right). Data represent mean ± SEM of 5 independent experiments, each performed in triplicate. Statistical significance was determined using a one-way ANOVA followed by Dunnett's multiple comparisons test against the WT group on the unnormalized raw data (left panel). Except for the negative control (**$P < 0.01$), differences in expression of all tested variants compared to WT were not significant, while the exact P value is shown for the D336A variant (~88% of WT, $P = 0.0614$).

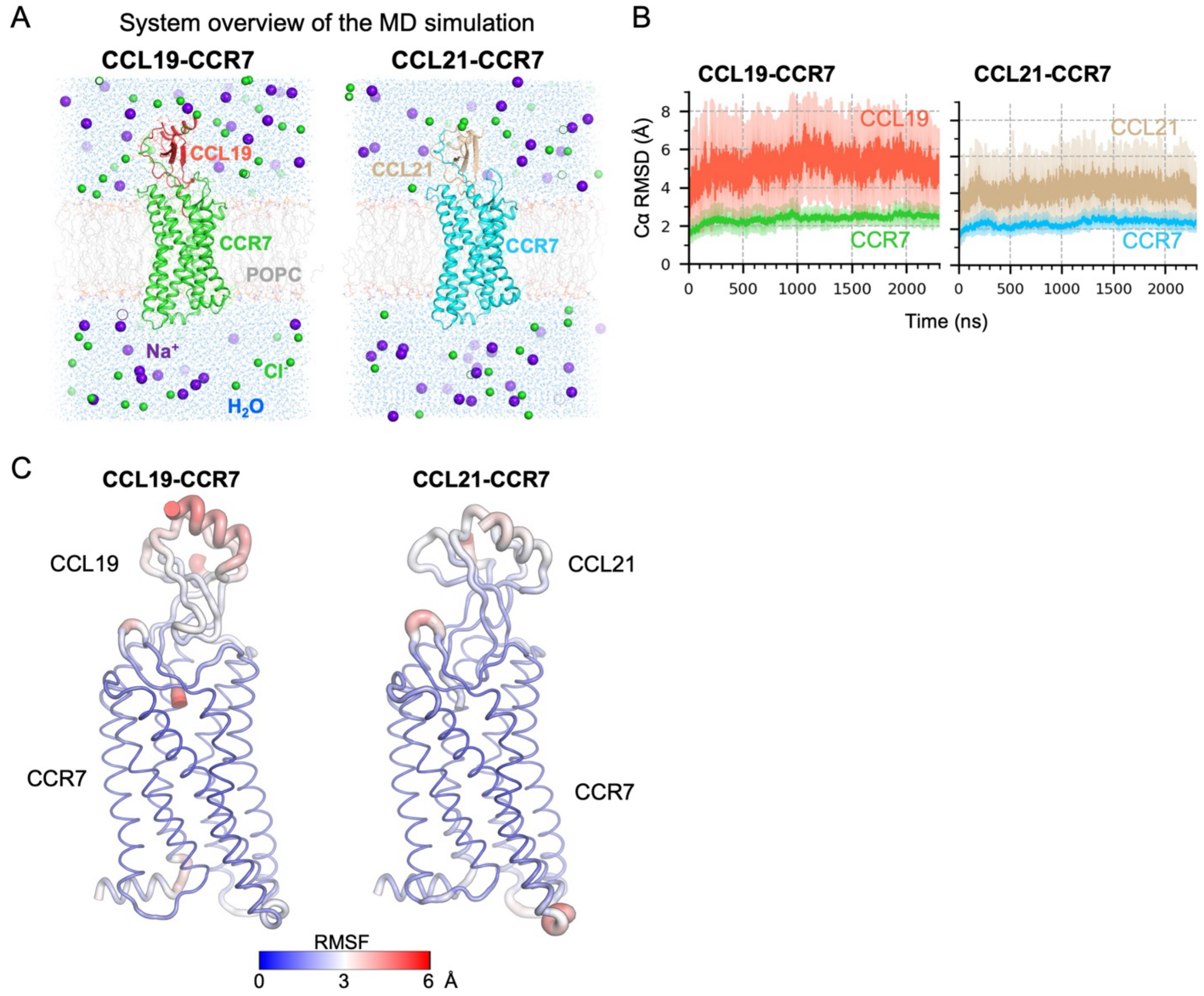


**Fig. S9 | Overview of the MD simulations.**

(A) Overview of the MD simulation system. The $G_i$ proteins and scFvs were removed from the cryo-EM models. The ligand-receptor pairs (shown as cartoons) were embedded in the POPC bilayers (shown as red/white lines), and were solvated with TIP3P water molecules (small blue dots) and 150 mM $Na^+$ (purple spheres) and $Cl^-$ (green spheres) ions. For each system, six independent 2.3 µs MD runs (replicas) were performed. (B) After aligning the trajectories by the CCR7 Cα atoms (see Methods), the RMSD time courses were measured for CCR7 and CCL19/21, respectively, without further alignment. The mean ± standard deviation of RMSD over the six replicas is shown in the graphs. (C) The average structures and the fluctuations calculated from the final 1.3 µs (1-2.3 µs) of the six replicas. The Cα traces are shown as tubes with thicknesses and colors to represent the root mean squared fluctuation (RMSF).

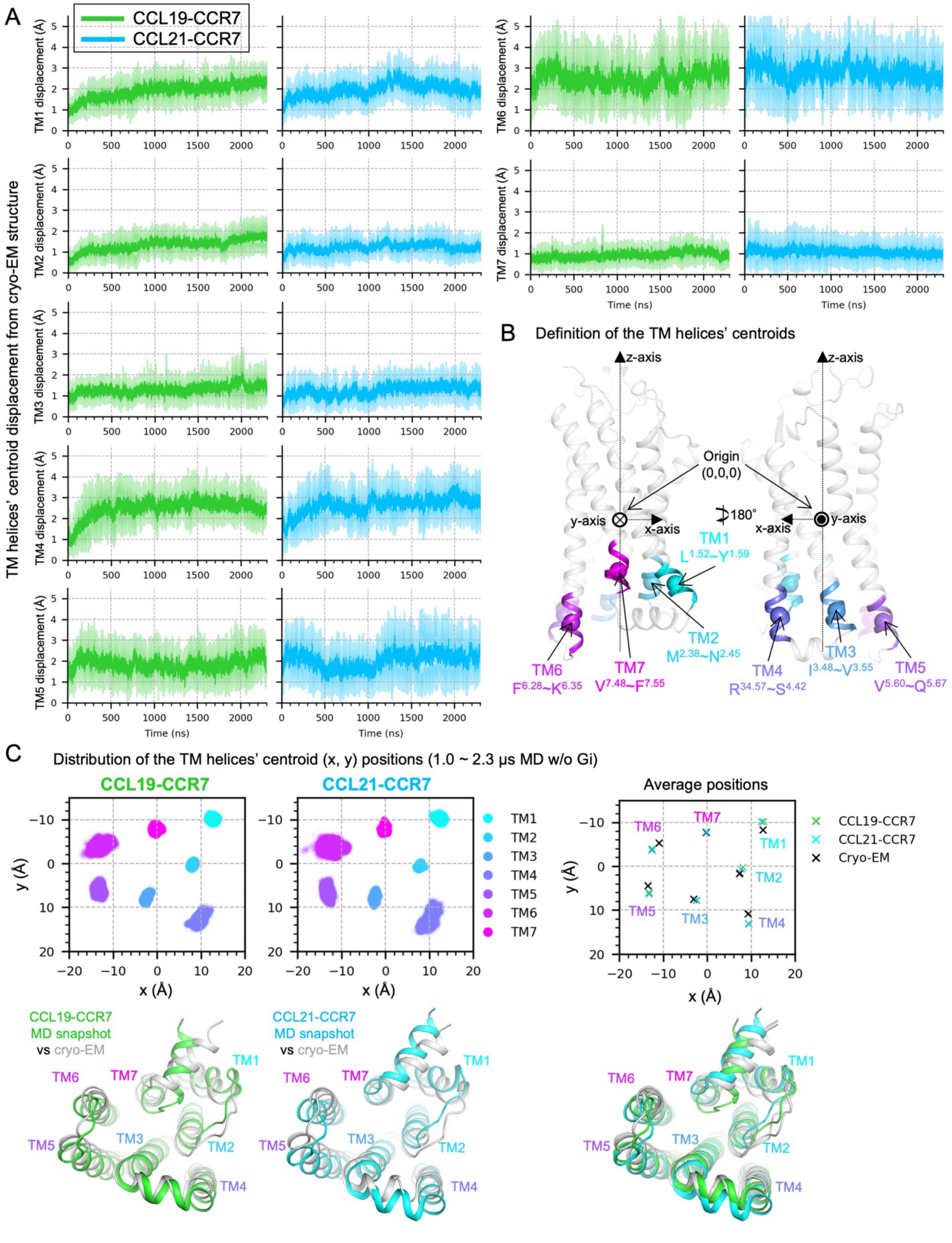


*Fig. S10 caption on the next page.*

**Fig. S10 | Measurement of the displacements of the TM helices in the intracellular side during the MD simulations.**

(A) Displacements of the TM helices' centroids (as defined in panel B) from the cryo-EM models over time. The mean ± standard deviation of the displacements over the six replicas is shown in the graphs. (B) Definition of the TM helices' centroids in the intracellular side. Trajectories were aligned to the cryo-EM model, which was transformed so the (x, y) origin is the protein center, the z origin is the membrane center, and the z-axis is perpendicular to the membrane. Each TM helix centroid was then defined as the centroid of Cα atoms of the two helical turns in the intracellular side (TM1: $L^{1.52}$ - $Y^{1.59}$, TM2: $M^{2.38}$ - $N^{2.45}$, TM3: $I^{3.48}$ - $V^{3.55}$, TM4: $R^{34.57}$ - $S^{4.42}$, TM5: $V^{5.60}$ - $Q^{5.67}$, TM6: $F^{6.28}$ - $K^{6.35}$, TM7: $V^{7.48}$ – $F^{7.55}$). (C) Top left and middle: Scatter plots of the (x, y) coordinates of the TM helices' centroids as defined in panel B. The coordinates from all MD frames except the first 1 µs were plotted. Top right: The average position of the TM helices' centroids is plotted, along with those of the cryo-EM models. Note that the average centroid positions for both complexes are essentially identical to each other. This indicates that the two ligands do not differentiate the intracellular TM positions after $G_i$ dissociation. Bottom left and middle: the MD snapshots and the corresponding cryo-EM models are superposed. The viewing direction is the same as the scatter plots (top row). Bottom right: The snapshots shown in the bottom left and middle are all superposed for comparison.

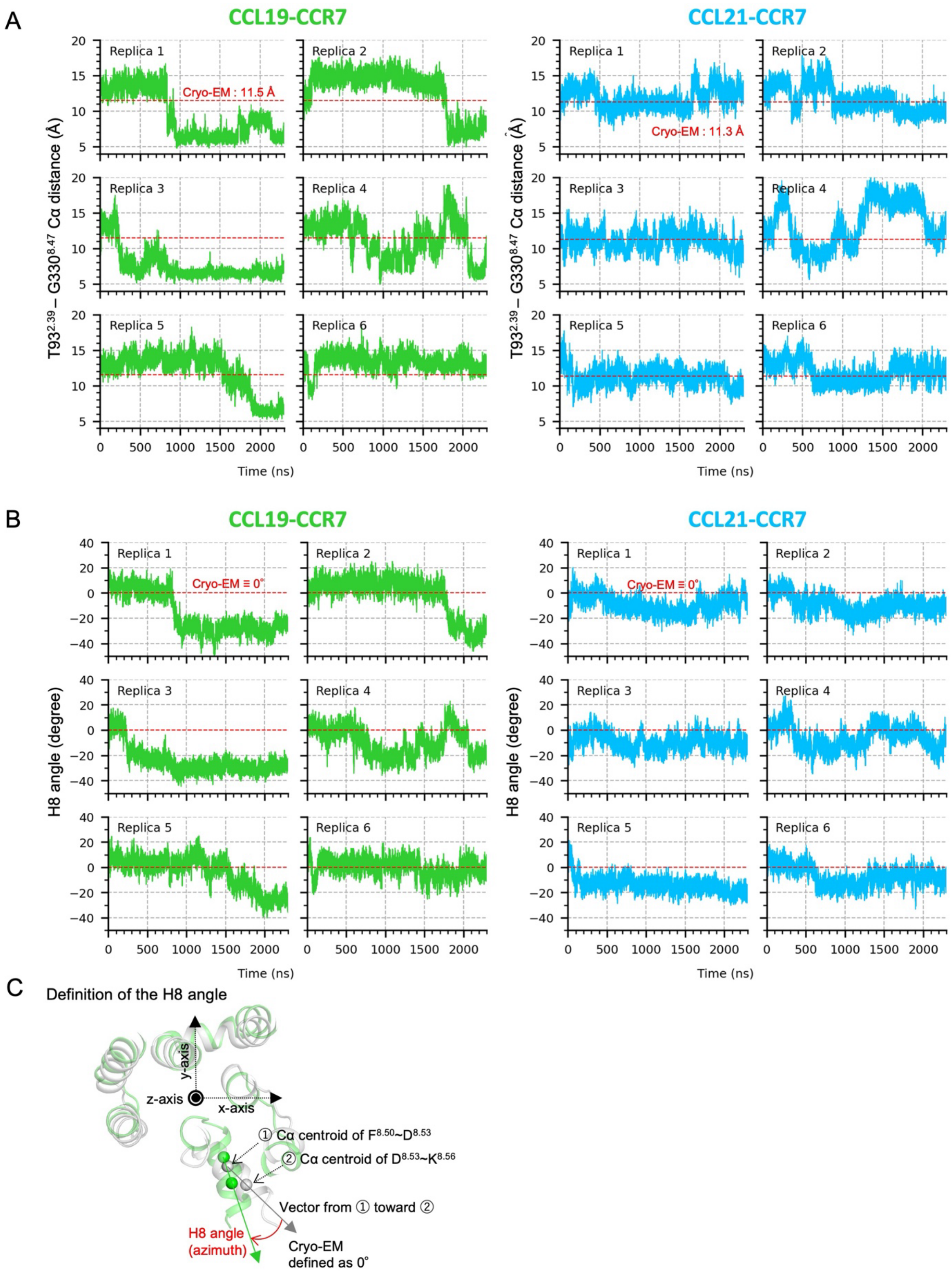


*Fig. S11 caption on the next page.*

**Fig. S11 | Analysis of the H8 orientation during the MD simulations.**

(A) Time courses of the Cα distance between T93$^{2.39}$ (TM2) and G330$^{8.47}$ (TM7-H8 loop). The raw data from six independent simulation replicas (Replica 1-6) are shown for CCL19-CCR7 (green) and CCL21-CCR7 (cyan). The corresponding distances in the cryo-EM structures (CCL19-bound: 11.5 Å, CCL21-bound: 11.3 Å) are indicated by red dashed lines. (B) Time courses of the H8 helix angle, as defined in (C). The raw data from the six replicas are shown for CCL19-CCR7 (green) and CCL21-CCR7 (cyan). The cryo-EM structure angle is defined as 0° (red dashed line). The T93$^{2.39}$ – G330$^{8.47}$ Cα distance and H8 angle data from the final 1.3 µs (1-2.3 µs) of these time courses were used to generate the probability density plots shown in Fig. 5A. (C) Definition of the H8 angle. After the trajectories were aligned as in Fig. S10C, for each MD frame, the vector from the Cα centroid of residues F$^{8.50}$ to D$^{8.53}$ (closer to the origin) toward the Cα centroid of residues D$^{8.53}$ to K$^{8.56}$ (further from the origin) was calculated, then projected onto the x-y plane (membrane plane). The angle between the projected vectors of the MD-frame and the cryo-EM model in the x-y plane was defined as the H8 angle (azimuth).

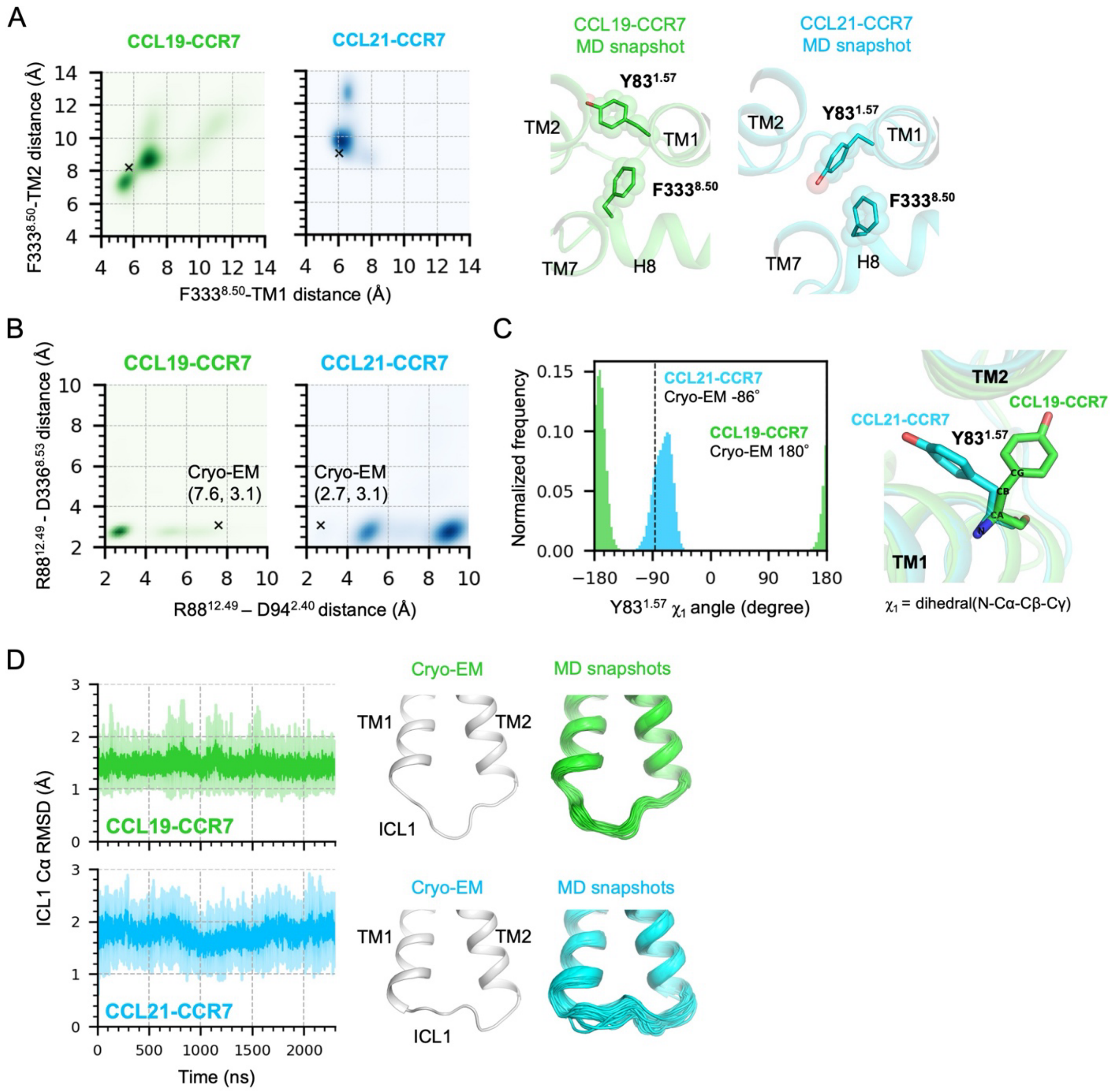


**Fig. S12 | Analysis of the H8 interaction with TM1, TM2 and ICL1 during the MD simulations.**

(A) Structural basis of the distinct H8 orientation between CCL19-bound CCR7 and CCL21-bound CCR7 in the $G_i$-free MD simulations. The closer positioning of H8 towards ICL1 and TM2 in CCL19-bound CCR7 is facilitated by the closer positioning of F333$^{8.50}$ towards TM1 and TM2, which is enabled by the outward-facing Y83$^{1.57}$. Left: The probability density as a function of the distance between F333$^{8.50}$ and the TM1 centroid or the TM2 centroid. See Fig. S10B for the definition of the TM centroids. Middle and right: A close-up view of H8, TM1, ICL1, and TM2 in the MD snapshots of CCL19-bound CCR7 (middle) and CCL21-bound CCR7 (right), with F333$^{8.50}$ and Y83$^{1.57}$

shown as sticks and spheres. (B) The probability density as a function of the R88$^{12.49}$ – D94$^{2.40}$ Cα distance and the R88$^{12.49}$ – D336$^{8.53}$ Cα distance, calculated from the $G_i$-free MD simulations (six independent 2.3 µs MD each. The first 1 µs was excluded from the analysis as a relaxation period). The distances from the cryo-EM models are indicated by black 'x' markers and text. (C) Left: The histogram of the χ1 dihedral angle of Y83$^{1.57}$ during the MD simulations. The same rotamers as in the cryo-EM models were maintained during the entire MD simulations. Right: The MD snapshots are superposed. Y83$^{1.57}$ is shown as sticks. The definition of the χ1 angle is indicated. (D) Left: After aligning the trajectories by the ICL1-flanking residues of TM1 and TM2 (see Methods), the RMSD of the ICL1 Cα atoms against the cryo-EM models was measured without further alignment. The mean ± SD over the six replicas is shown in the graphs. Middle and right: The cryo-EM models and the MD snapshots (every 100 ns in 1.0 ~ 2.3 µs from six replicas, i.e., 84 models are superposed) are shown in cartoons. The low RMSD values indicate that the distinct ICL1 conformations, which correspond to the $G_i$-bound states observed in the cryo-EM structures, are stably maintained in the $G_i$-free simulations.

**Table S1. Data collection and refinement statistics of the cryo-EM analysis.**

| | CCL19-CCR7-$G_i$-scFv16 (EMDB-66874, PDB 9XHH) | CCL21-CCR7-$G_i$-scFv16 (EMDB-66875, PDB 9XHI) |
|---|---|---|
| Data collection and processing | | |
| Microscope | JEM-Z320FHC (JEOL) | |
| Calibrated magnification | 63,291x | |
| Voltage (kV) | 300 | |
| Electron exposure (e/Å$^2$) | 50 | |
| Defocus range (μm) | -1.0 to -1.8 | |
| Pixel size (Å) | 0.79 | |
| Symmetry imposed | C1 | C1 |
| Micrographs used (no.) | 20,142 | 5,968 |
| "Initial" particle images (no.) | 3,374,958 | 1,537,777 |
| Final particle images (no.) | 256,195 | 195,011 |
| Map resolution (Å)<br>FSC threshold | 3.0<br>0.143 | 3.2<br>0.143 |
| Refinement | | |
| Initial model used | AlphaFold3 | AlphaFold3 |
| Model resolution (Å)<br>FSC threshold | 3.3<br>0.5 | 3.4<br>0.5 |
| Map sharpening method | uniform (-60 Å$^2$) | uniform (-60 Å$^2$) |
| Model composition<br>Non-hydrogen atoms<br>Protein residues | <br>9,463<br>1,216 | <br>9,449<br>1,213 |
| *B* factor (Å$^2$)<br>Protein<br>Ligand | <br>94.62<br>61.55 (CLR) | <br>90.30<br>48.93 (CLR) |
| R.m.s. deviations<br>Bond lengths (Å)<br>Bond angles (°) | <br>0.004<br>0.664 | <br>0.003<br>0.619 |
| Validation<br>MolProbity score<br>Clashscore<br>Poor rotamers (%) | <br>1.68<br>7.43<br>0 | <br>1.69<br>7.75<br>0 |
| Ramachandran plot<br>Favored (%)<br>Allowed (%)<br>Disallowed (%) | <br>95.99<br>4.01<br>0 | <br>96.15<br>3.85<br>0 |
| EMRinger score | 2.67 | 1.93 |
